# General Functional Connectivity: shared features of resting-state and task fMRI drive reliable and heritable individual differences in functional brain networks

**DOI:** 10.1101/330530

**Authors:** Maxwell L. Elliott, Annchen R. Knodt, Megan Cooke, M. Justin Kim, Tracy R. Melzer, Ross Keenan, David Ireland, Sandhya Ramrakha, Richie Poulton, Avshalom Caspi, Terrie E. Moffitt, Ahmad R. Hariri

## Abstract

Intrinsic connectivity, measured using resting-state fMRI, has emerged as a fundamental tool in the study of the human brain. However, due to practical limitations, many studies do not collect enough resting-state data to generate reliable measures of intrinsic connectivity necessary for studying individual differences. Here we present general functional connectivity (GFC) as a method for leveraging shared features across resting-state and task fMRI and demonstrate in the Human Connectome Project and the Dunedin Study that GFC offers better test-retest reliability than intrinsic connectivity estimated from the same amount of resting-state data alone. Furthermore, at equivalent scan lengths, GFC displays higher heritability on average than resting-state functional connectivity. We also show that predictions of cognitive ability from GFC generalize across datasets, performing as well or better than resting-state or task data alone. Collectively, our work suggests that GFC can improve the reliability of intrinsic connectivity estimates in existing datasets and, subsequently, the opportunity to identify meaningful correlates of individual differences in behavior. Given that task and resting-state data are often collected together, many researchers can immediately derive more reliable measures of intrinsic connectivity through the adoption of GFC rather than solely using resting-state data. Moreover, by better capturing heritable variation in intrinsic connectivity, GFC represents a novel endophenotype with broad applications in clinical neuroscience and biomarker discovery.

## Introduction

Functional magnetic resonance imaging (fMRI) has proven invaluable for identifying the neural architecture of human behavior and cognition (Betti et al., 2013; Fox et al., 2007; Huth et al., 2016). Accordingly, fMRI has been widely adopted in myriad studies seeking to deepen our understanding of the human brain in both health and disease (Cole et al., 2011; Power et al., 2013; Satterthwaite et al., 2016). Recently, fMRI studies have expanded in both scale and scope, often collecting data in thousands of individuals in an effort to adequately power the search for neural correlates of complex human traits and predictive biomarkers for mental illness (Casey et al., 2018; Elliott et al., 2018; Miller et al., 2016; Swartz et al., 2015; Thompson et al., 2014). In this context, most studies have focused on the acquisition of resting-state functional MRI to map the intrinsic connectivity of neural networks. This choice is often prompted by three considerations. First, intrinsic connectivity networks appear to be more heritable than task-elicited activity (Elliott et al., 2017; Ge et al., 2017; Winkler et al., 2010). Second, resting-state data are relatively easy to collect from informative populations of children, the elderly, and mentally ill patients (Fox, 2010; Greicius, 2008; Shehzad et al., 2009). Third, intrinsic connectivity plays a primary role in shaping task-based brain activity and associated behaviors (Cole et al., 2016, 2014; Krienen et al., 2014; Tavor et al., 2016).

While analyses of resting-state data have revealed insights about average, group-level features of the human brain (Buckner et al., 2008; Power et al., 2011; Yeo et al., 2011), progress has lagged in identifying individualized signatures, which are critical if ongoing large-scale studies are to be successful in the search for risk and disease biomarkers. This lack of progress partially reflects the generally poor test-retest reliability of intrinsic connectivity estimates in many resting-state datasets (Birn et al., 2013; Guo et al., 2012). The reliability of intrinsic connectivity measures must be improved, as unreliability limits the ability to predict individual differences that could become clinical biomarkers (Nunnally Jr., 1970; Vul et al., 2009). In this paper we propose general functional connectivity (GFC), a simple method for combining resting-state and task fMRI data to improve the reliability of intrinsic connectivity.

Historically researchers have collected 5-10 minutes of resting-state data in response to both published recommendations (Shehzad et al., 2009; Van Dijk et al., 2010) and practical limitations (e.g., scan time, cost and subject stamina). While such data may be adequate for detecting patterns of intrinsic connectivity common across individuals, more than 25 minutes of data are needed to reliably detect individual differences in these same patterns (Anderson et al., 2011; Hacker et al., 2013; Laumann et al., 2015). This suggests that intrinsic connectivity may simply be a noisy measure of stable individual differences and that lengthier measurement is needed to identify the “true” signal (Braga and Buckner, 2017; Gordon et al., 2017; Gratton et al., 2018; Mueller et al., 2013). However, due to the high cost of scan time and the limited ability of many individuals including children, the elderly, and the mentally ill to lie still without falling asleep for tens of minutes of scanning, this level of protracted measurement is simply not feasible for a majority of studies (Power et al., 2012; Satterthwaite et al., 2013; Tagliazucchi and Laufs, 2014). To illustrate the point, two recent meta-analyses of resting-state intrinsic connectivity in depression (Kaiser et al., 2015) and schizophrenia (Dong et al., 2018) revealed average resting-state scan times of 6.53 minutes (k = 23 studies) and 6.24 minutes (k = 36 studies), respectively. It is likely that intrinsic connectivity based on these brief scans has low reliability, reducing the ability to discover individual differences and related biomarkers. If measures of intrinsic connectivity are to realize their potential as a clinical tool (Fox, 2010; Matthews et al., 2006) and contribute to personalized medicine (Collins and Varmus, 2015; Hamburg and Collins, 2010), there must first be substantial improvement in their reliability, and this should be ideally achieved within the practical limitations of study design and participant burden.

Although few studies collect enough resting-state data to achieve high levels of reliability, many collect multiple task scans in addition to a resting-state scan. While it has been implicitly assumed that task and resting-state scans are two separate measures of brain function, to be analyzed and studied independently, a growing body of evidence suggests that the intrinsic connectivity measured by each may have substantial overlap, sharing over 80% of the same variance (Cole et al., 2014; Geerligs et al., 2015; Tavor et al., 2016). In fact, it has long been known that intrinsic networks extracted from task scans are quite similar to those extracted from resting-state scans, and task scans have been used for intrinsic connectivity analyses when resting-state data are absent (Arfanakis et al., 2000; Fair et al., 2007; Fox et al., 2006; Smith et al., 2009). Further, mental operations that occur during task states have been linked to resting-state functional networks (Bzdok et al., 2016; Leech et al., 2012), and task scans have been shown to accentuate individual differences in network dynamics (Finn et al., 2017; Greene et al., 2018; Satterthwaite et al., 2018). Despite this evidence for a complementary relationship between resting-state and task fMRI, they are typically investigated separately. Even when researchers have collected both task and resting-state fMRI in the same subjects, analyses of intrinsic connectivity are almost always performed on a single short resting-state scan despite the low test-retest reliability of short scan data. If it could be demonstrated that the reliability of intrinsic connectivity can be improved by adding task to resting-state fMRI, many researchers could immediately benefit by adopting this approach in data they have already collected, and also in designing future studies. The field of individual-differences neuroscience would also benefit from the resulting increase in reliability, replicability, and power that would follow the widespread adoption of combining task and resting-state fMRI.

Here we hypothesized that measures of intrinsic connectivity will be more reliable when task and resting-state scans are combined into a longer dataset (i.e., GFC) compared to short resting-state scans alone. Additionally, we hypothesized that this boost in reliability with GFC will increase the amount of heritable variation detectable in intrinsic connectivity, because the detection of heritability requires reliable measurement. Finally, we hypothesized that GFC would measure a common backbone of intrinsic connectivity common across task and resting-state data allowing for better out-of-sample prediction of individual differences. We examined two existing datasets in 3 different ways to test these hypotheses. First, we investigated the test-retest reliability of different combinations of task and resting-state data in both the Human Connectome Project (HCP (Van Essen et al., 2013)), which uses state-of-the-art fMRI hardware and acquisition parameters in a highly educated, healthy sample, and in the Dunedin Study (Poulton et al., 2015), which uses more common fMRI hardware and acquisition parameters in a population-representative birth cohort. Second, we estimated the heritability of GFC and resting-state functional connectivity (RSFC) by leveraging the twin sampling design of the HCP. Third, we investigated the ability of GFC to measure generalizable individual differences in cognitive ability across samples and across task states by testing out-of-sample prediction of cognitive ability in the HCP and Dunedin Study samples.

## Materials and Methods

### Datasets

#### Human Connectome Project

This is a publicly available data set that includes 1206 participants with extensive MRI and behavioral measurement (Van Essen et al., 2013). In addition, 45 of these subjects completed the entire scan protocol a second time (referred to hereafter as “test-retest sample”). All participants were free of current psychiatric or neurologic illness and were 25-35 years of age. In all analyses, subjects were excluded if they had truncated scans, less than 40 minutes of combined task data, or less than 40 minutes of resting-state data after censoring. Much of the HCP dataset consists of twins and family members. We exploited this feature of the design for conducting heritability analyses. For these analyses, 943 participants from 420 families met inclusion criteria (144 monozygotic families, 85 dizygotic families, and 191 full sibling families). To avoid bias due to family confounding in the prediction analyses, only one individual per family was retained (the family member with the least motion during scanning). This resulted in 298 subjects left to be used in our heritability analyses. Eight subjects were removed from our test-retest analyses because they had at least one fMRI scan with truncated or missing data. Four subjects were removed because they did not have sufficient data (i.e., > 40 minutes of rest and > 40 minutes of task) after censoring. Therefore, the test-retest reliability analyses included data from 33 participants.

The acquisition parameters and minimal preprocessing of these data have been described extensively elsewhere (Glasser et al., 2013). In our analyses, we used the minimally preprocessed data in volumetric Montreal Neurological Institute (MNI) space (“fMRIVolume” pipeline). Briefly, participants underwent extensive MRI measurement that included T1 and T2 weighted structural imaging, diffusion tensor imaging, and nearly two hours of resting-state and task fMRI. One hour of resting-state fMRI was collected on each participant in four 15-minute scans (1200 time-points each) split-up into two scanning sessions over two days. In each scan session the two resting-state scans were followed by task fMRI (Smith et al., 2013). Across the two sessions, each participant completed seven fMRI tasks described extensively elsewhere (Barch et al., 2013). Briefly, tasks were designed to identify functionally relevant “nodes” in the brain and included working memory (810 timepoints, 10:02 minutes), gambling (506 timepoints, 6:24 minutes), motor function (568 timepoints, 7:06 minutes), language (632 timepoints, 7:54 minutes), social cognition (548 timepoints, 6:54 minutes), relational processing (464 timepoints, 6:52 minutes) and emotional processing (352 timepoints, 4:32 minutes). Altogether, 4800 timepoints totaling 60 minutes of resting-state fMRI and 3880 timepoints totaling 48:30 minutes of task fMRI were collected on each participant.

#### Dunedin Study

This is a longitudinal investigation of health and behavior in a complete birth cohort of 1,037 individuals (91% of eligible births; 52% male) born between April 1972 and March 1973 in Dunedin, New Zealand (NZ), and eligible based on residence in the province and participation in the first assessment at age three. The cohort represents the full range of socioeconomic status on NZ’s South Island and matches the NZ National Health and Nutrition Survey on key health indicators (e.g., BMI, smoking, GP visits) (Poulton et al., 2015). The cohort is primarily white; fewer than 7% self-identify as having non-Caucasian ancestry, matching the South Island (Poulton et al., 2015). Assessments were carried out at birth and ages 3, 5, 7, 9, 11, 13, 15, 18, 21, 26, 32, and 38 years, when 95% of the 1,007 study members still alive took part. Neuroimaging data collection is ongoing in participants who are now aged 45 years. We currently have collected completed neuroimaging data for 756 study members. Data were excluded if the study member did not complete the rest scan and all four task scans or if they did not have sufficient data after censoring and bandpass filtering was applied to the resting-state and each task scan separately (see below for censoring details). Data for 591 subjects survived these fMRI exclusion criteria and were included in analyses. Additionally, 20 study members completed the entire scan protocol a second time (mean days between scans = 79). Of these one failed to have sufficient degrees of freedom after censoring and bandpass filtering. Therefore, the test-retest reliability analyses included data from 19 participants.

Each participant was scanned using a Siemens Skyra 3T scanner equipped with a 64-channel head/neck coil at the Pacific Radiology imaging center in Dunedin, New Zealand. High resolution structural images were obtained using a T1-weighted MP-RAGE sequence with the following parameters: TR = 2400 ms; TE = 1.98 ms; 208 sagittal slices; flip angle, 9°; FOV, 224 mm; matrix =256×256; slice thickness = 0.9 mm with no gap (voxel size 0.9×0.875×0.875 mm); and total scan time = 6:52 minutes. Functional MRI was collected during resting-state and four tasks with a series of 72 interleaved axial T2-weighted functional slices acquired using a 3-fold multi-band accelerated echo planar imaging sequence with the following parameters: TR = 2000 ms, TE = 27 ms, flip angle = 90°, field-of-view = 200 mm, voxel size = 2mm isotropic, slice thickness = 2 mm without gap.

8:16 minutes (248 TRs) of resting-state fMRI was collected immediately before the four task fMRI scans. During the resting-state scan participants were instructed to stay awake with their eyes open while looking at a grey screen. Participants completed an emotion processing task (6:40 minutes, 200 TRs), a color Stroop task (6:58 minutes, 209 TRs), a monetary incentive delay (MID) task (7:44 minutes, 232 TRs) and an episodic memory task (5:44 minutes, 172 TRs). All four tasks are described in detail in the supplement.

### JMRI pre-processing

Minimal preprocessing was first applied to all data. For the HCP dataset this was done with the HCP minimal preprocessing pipeline (Glasser et al., 2013) and included correction for B0 distortion, realignment to correct for motion, registration to the participant’s structural scan, normalization to the 4D mean, brain masking, and non-linear warping to MNI space.

Minimal preprocessing steps were applied to the Dunedin Study dataset using custom processing scripts. Anatomical images were skull-stripped, intensity-normalized, and nonlinearly warped to a study-specific average template in the standard MNI stereotactic space using the ANTs SyN registration algorithm (Avants et al., 2008; Klein et al., 2009). Time-series images were despiked, slice-time-corrected, realigned to the first volume in the time-series to correct for head motion using AFNI tools (Cox, 1996), coregistered to the anatomical image using FSL’s Boundary Based Registration (Greve and Fischl, 2009), spatially normalized into MNI space using the non-linear ANTs SyN warp from the anatomical image, and resampled to 2mm isotropic voxels. Dunedin Study images were additionally corrected for B0 distortions.

Time-series images for each dataset were further processed to limit the influence of motion and other artifacts. Voxel-wise signal intensities were scaled to yield a time-series mean of 100 for each voxel. Motion regressors were created using each participant’s 6 motion correction parameters (3 rotation and 3 translation) and their first derivatives (Jo et al., 2013; Satterthwaite et al., 2013) yielding 12 motion regressors. Five components from white matter and cerebrospinal fluid were extracted using CompCorr (Behzadi et al., 2007) and used as nuisance regressors, along with the mean global signal. In the Dunedin Study dataset, images were bandpass filtered to retain frequencies between .008 and .1 Hz. In the HCP dataset, images only underwent highpass filtering with a cutoff of .008 Hz. High frequency signals were retained because removing high frequency signals would have resulted in excessive loss of degrees of freedom due to the very low TR (.75 seconds) (Bright et al., 2017; Caballero-Gaudes and Reynolds, 2017). In the HCP dataset, we followed the empirically derived thresholds of .39 mm frame-wise displacement or 4.9 units above the median DVARS as recommended (Burgess et al., 2016). In the Dunedin Study dataset, we investigated a range of framewise-displacement cutoffs using QC-RSFC plots in order to derive the optimal threshold for removing motion artifact as recommended (Power et al., 2014). This investigation led to thresholds of 0.35 mm framewise-displacement and 1.55 standardized DVARS. In both datasets, nuisance regression, bandpass filtering, censoring, global-signal regression, and smoothing of 6mm FWHM for each time-series were performed in a single processing step using AFNI’s 3dTproject. Identical time-series processing was applied to resting-state and task time-series data with one exception.

To remove functional connectivity predominantly driven by task-evoked coactivation, signal due to task structure was added as an additional nuisance covariate to all task scans and removed from their time-series (Fair et al., 2007) (see supplemental for details on task modeling and regression). However, we examined the impact of task structure on our estimates of intrinsic connectivity by conducting parallel analyses without the regression of task structure. While task regression had a small but consistent effect of lowering ICCs for intrinsic connectivity estimates (see supplemental Table S1), the ability to predict cognitive ability was slightly higher when task regression was applied (see supplemental Table S2). Thus, we focus on analyses with task regression in our primary results.

### Effects of Global Signal Regression

Global signal regression (GSR) was performed to adopt a conservative approach to motion artifact reduction and used in our main analyses (Power et al., 2018). However, because use of GSR is still debated (Murphy and Fox, 2017), we also preprocessed our data with parallel preprocessing methods without GSR (see supplemental figure S1).

### Functional Connectivity processing

We investigated whole-brain intrinsic connectivity using two parcellation schemes. In the reliability and prediction analyses (described below), a 264-region parcellation was utilized. This parcellation was derived in a large independent dataset (Power et al., 2011). BOLD data were averaged within 5mm spheres surrounding each of the 264 coordinates in the parcellation. In the heritability analyses we used a smaller 44-brain-region parcellation scheme due to computational constraints imposed by the time needed to fit heritability models at each edge independently (described below). This parcellation was derived from the 7-network parcellation scheme described in (Yeo et al., 2011). These 7 networks were broken up into 44 spatially contiguous regions across the two hemispheres. Average time series were extracted from all 44 regions. For both parcellations the average time-series was extracted independently from each scan session in every dataset. This allowed the time-series to be flexibly concatenated and recombined. Correlation matrices were derived from these time-series using Pearson correlation, resulting in 34,716 edges in the Power et al. parcellation and 946 edges in the Yeo et al. parcellation.

### Test-retest reliability

Intraclass correlation (ICC) was used to quantify the test-retest reliability of intrinsic connectivity measures in the HCP and Dunedin Study test-retest datasets. ICC (3,1) was used in all analyses (Chen et al., 2018). In the HCP dataset, the influence of the amount of data, as well as type of data (resting-state or task) on ICC was investigated. In the resting-state analyses, intrinsic connectivity matrices were calculated across a range of scan lengths (5, 10, 20, 30, and 40 minutes of post-censoring data) mirrored in the test and retest samples. ICCs were then calculated for each edge of this matrix, yielding 34716 (264*263/2) unique ICC values for each scan length. A more general definition of intrinsic connectivity was derived by including task data in the intrinsic connectivity matrices. GFC was formally investigated (Figures 1, 2 and 3) by constructing datasets from a combination of task and resting-state data over a variety of scan lengths (5, 10, 20, 30 and 40 minutes of post-censoring data). In scan lengths up to 25 minutes, an exactly equal amount of data from all scan types (1/8 of total scan length from each task and resting-state) was combined. After this point, equal amounts of each task could no longer be added together because the shortest task scan no longer had sufficient timepoints remaining in all participants. Above 25 minutes, timepoints were selected at random from the pool of remaining timepoints. In the Dunedin Study, two ICC matrices were constructed. The first was built from each participant’s single resting-state scan and the second from all task and resting-state scans to create the GFC matrix. Paired-sample t-tests were used to compare mean differences in reliability between ICC matrices constructed from traditional resting-state functional connectivity and GFC.

**Figure 1.**
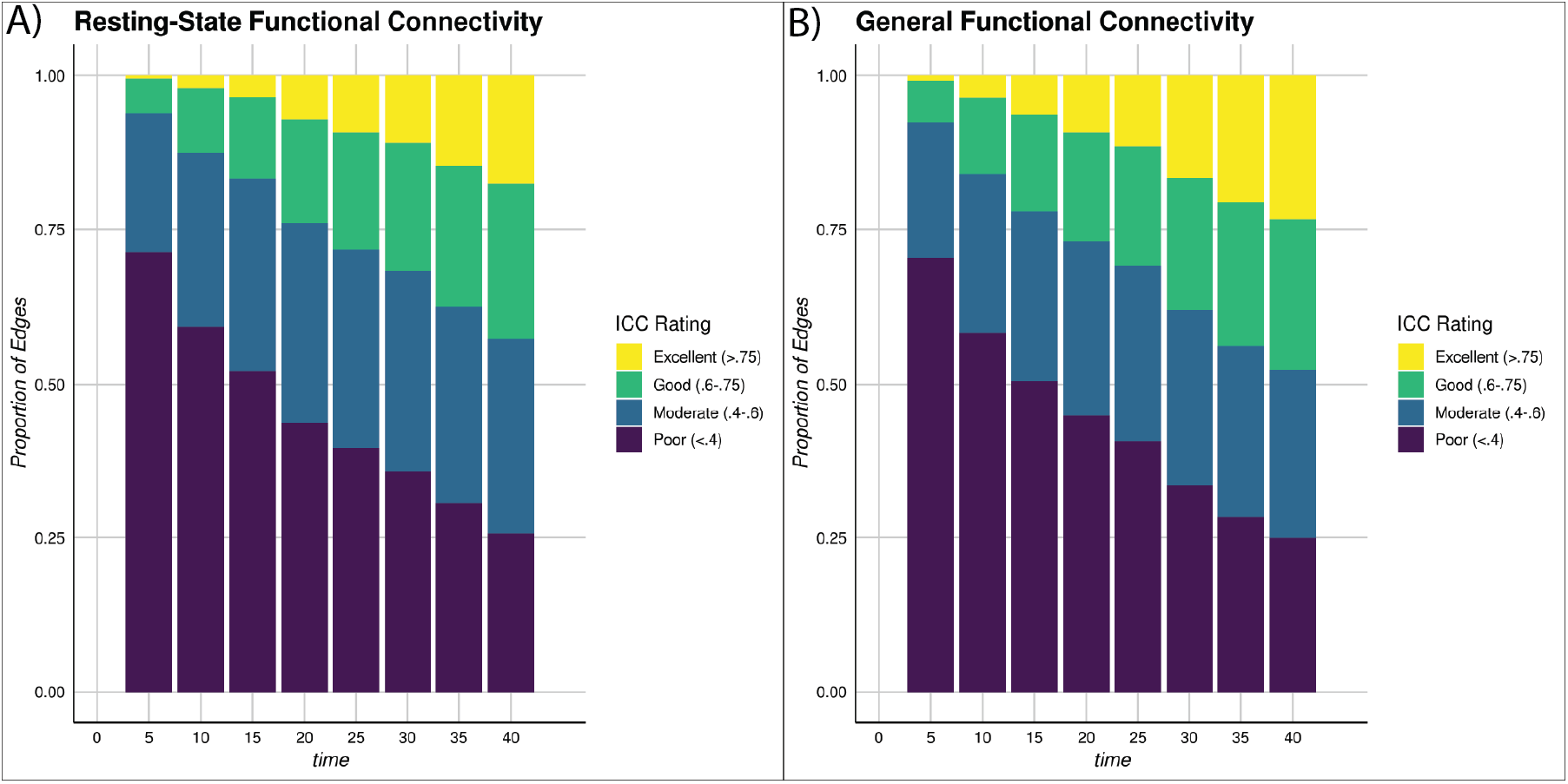
Test-retest reliability of individual differences in global intrinsic network connectivity increases as the amount of data used to estimate either RSFC (A) or GFC (B) increases. Stacked bar plots from the HCP dataset displaying the proportion of functional connections (i.e., edges) across neural networks as defined by (Power et al., 2011) that meet criteria for excellent, good, moderate, and poor reliability as indexed by ICCs.

**Figure 2.**
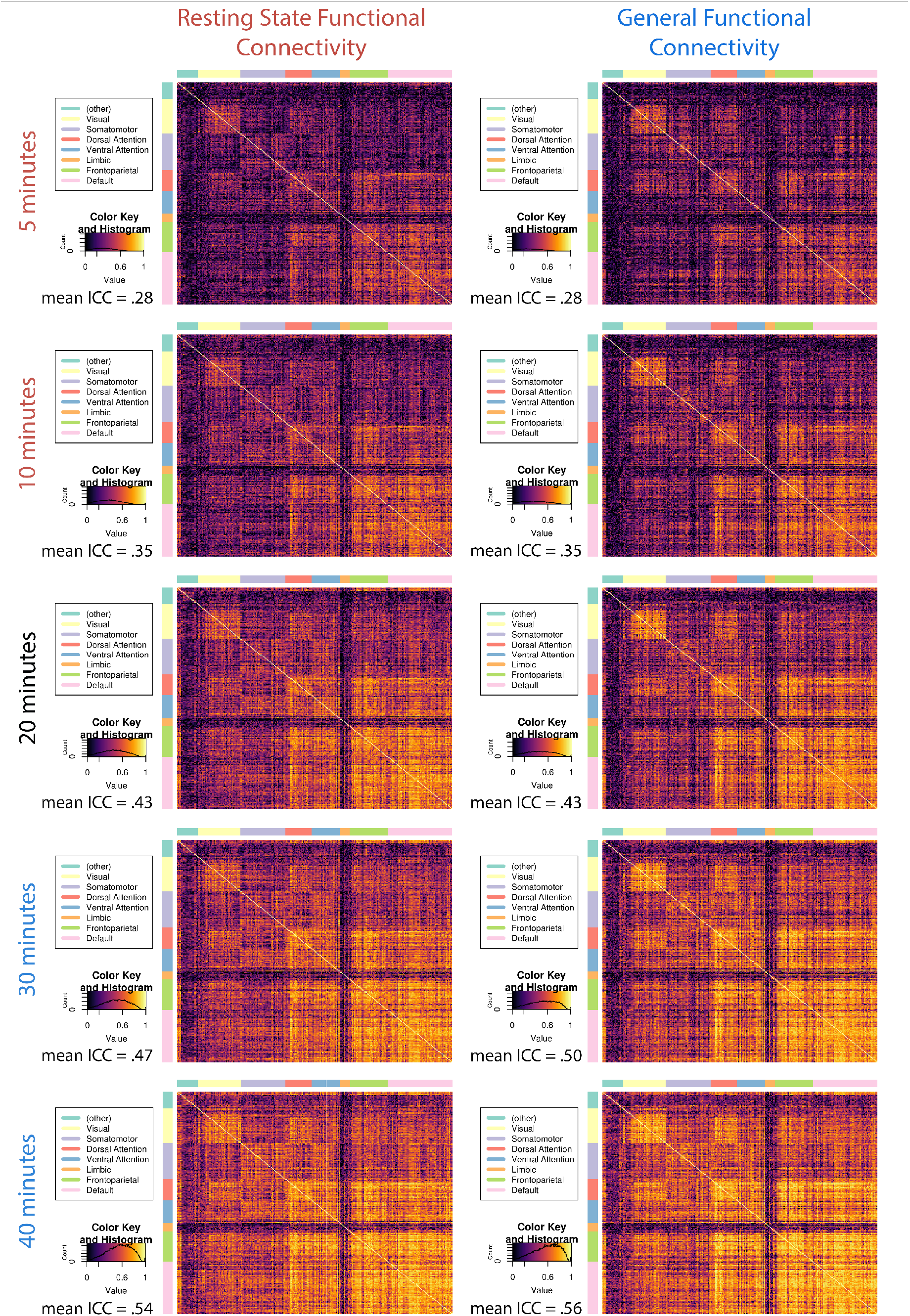
Test-retest reliability of the intrinsic connectivity of canonical neural networks derived from either RSFC or GFC from the HCP scales with the amount of data available for analysis. Intra-Class Correlation (ICC) matrices for RSFC (left panel) and GFC (right panel) demonstrate comparable gains in reliability with increasing amounts of data across common intrinsic networks (Power et al., 2011). 5 and 10 minutes are written in red because these are common scan lengths for resting-state scans. 30 and 40 minutes are written in blue because many researchers have collected this amount of fMRI data when resting-state and task scans are combined. To the bottom left of ICC matrices is the color key for the ICCs, with a histogram indicating the density of ICCs for the corresponding graph.

HCP fMRI data were collected in two phase-encoding directions (LR and RL). To minimize bias due to phase-encoding direction on our analyses, in each participant, for each task, timepoints were first selected from LR encoded scans until LR timepoints were exhausted. Subsequently, timepoints were selected from remaining RL-encoded scans. This ensured that all subjects had a roughly equivalent proportion of LR and RL timepoints within a given analysis. This also ensured roughly equivalent proportions of LR and RL phase encoding across test and retest correlation matrices when calculating ICCs. Selecting from LR first before RL had the additional benefit of mimicking shorter data-acquisition scans because timepoints were utilized in the order they were collected.

### Heritability

For each edge of the connectivity matrix, we ran an extended twin model (Posthuma and Boomsma, 2000), which is an extension of the classic twin design in that it allows for the modeling of additional family members of twins. In the HCP study we had data from the families of monozygotic and dizygotic twins as well as siblings in families without a twin pair. In this model the variance of the phenotype (in this case correlations between brain regions) is partitioned into four sources: additive genetic factors (A), shared environmental factors (C), twin-specific factors (T), and unique environmental factors and error (E) based on the correlations between monozygotic twins, dizygotic twins, and their siblings. Due to their small numbers, we eliminated half-siblings from the model (N = 27) and capped family size at four individuals (7 families affected, N = 13). This left us with 1,078 individuals from 443 families (148 monozygotic families, 92 dizygotic Z families, 203 full-sibling families). Of these, 943 participants from 420 families (144 monozygotic families, 85 dizygotic families and 191 full-sibling families) had full scans from all tasks and rest and had at least 40 minutes of task data and 40 minutes of rest data after censoring. Heritability analyses were performed using OpenMx (Boker et al., 2011; Neale et al., 2016) in the R statistical computing environment. Each model also contained parameters to adjust for the effects of sex and age on the mean. Confidence intervals were obtained on the A, C, T, and E estimates using the mxCI command in OpenMx which obtains profile likelihood confidence intervals (Pek and Wu, 2015). For each edge, an estimate was determined to be significant if the lower bound of the confidence interval was greater than zero.

### Connectome-basedpredictive modeling

The predictive utility of increasing reliability through our GFC measure was evaluated by testing the ability of the different intrinsic connectivity matrices to predict cognitive ability in the HCP and Dunedin Study datasets. Cognitive ability was measured in the HCP using the Raven’s Progressive Matrices (PMAT24_A_CR) (Raven, 1941). This measure was adopted because it is a proxy for cognitive ability that has been shown to be predicted by intrinsic connectivity in the HCP (Dubois et al., 2018; Finn et al., 2015; Noble et al., 2017). The WAIS-IV, a well-established and validated measure of individual differences in cognitive ability (Weschler, 2008), was used in the Dunedin Study.

Cognitive ability was predicted from intrinsic connectivity data using connectome-based predictive modeling (CPM; (Shen et al., 2017)). This framework provides a general method to predict any measure from intrinsic connectivity matrices. In this approach, predictors are filtered by selecting edges that have a p < .01 correlation with the measure of interest. Three linear regression predictive models are then built, one from the positive features (edges positively correlated with the phenotype of interest), one from the negative features (edges negatively correlated with the phenotype of interest), and one from the combination of positive and negative features (Shen et al., 2017). Here we report results from the combined model (see supplemental tables S3, S4 and S5 for results from all models). In our CPM analyses, we used both within-sample and out-of-sample prediction. Within-sample models were used to directly compare RSFC and GFC predictions of cognitive ability using a leave-one-out cross-validation scheme. Models were trained with all participants except one and used to predict the measure in the left-out participant. This was repeated until all participants had been left out.

Out-of-sample prediction is the gold standard to test the unbiased predictive utility of models (Whelan and Garavan, 2014; Yarkoni and Westfall, 2017) and was therefore used to test the generalizability of predictive models built with RSFC, GFC, and task data. To test the influence of task state, data aggregation and scan length on predictive utility, models were trained with RSFC, GFC, and individual tasks in the Dunedin Study dataset and then used to predict cognitive ability from RSFC, GFC, and individual tasks in the HCP dataset. The Dunedin Study and HCP have three parallel tasks that tap into similar behavioral circuits. The gambling task in the HCP and the MID task in Dunedin Study both involve reward processing. The emotional processing task in the HCP and the face matching task in the Dunedin Study both involve passive processing of emotional facial expressions. Lastly, the working memory task in the HCP and the Stroop task in the Dunedin Study both tap into executive control. We restricted our analyses to comparisons of RSFC, GFC, and these 3 parallel tasks. The abilities of RSFC and GFC was contrasted with that of these parallel tasks to predict cognitive ability within tasks. In both within-sample and out-of-sample tests the Spearman correlation between predicted and true scores was adopted as an unbiased effect size measure of predictive utility. Model predictions of cognitive ability were assessed for significance using a parametric test for significance of correlations. All p-values from correlations with cognitive ability were corrected for multiple comparisons using the false discovery rate (Benjamini and Hochberg, 1995). Differences in predictive utility between models were compared using Steiger’s z (Steiger, 1980). All confidence intervals for CPM prediction estimates were generated with bootstrap resampling, using AFNI’s 1dCorrelate tool.

## Results

### What is the test-retest reliability of GFC?

We used data from the HCP to evaluate the test-retest reliability of both intrinsic connectivity derived from resting-state scans alone and of GFC derived from combinations of task and resting-state scans. Test-retest reliability, as measured by ICC, ranges from 0-1, and is commonly classified according to the following cutoffs: 0 – 0.4 = poor, 0.4 – 0.6 = moderate, 0.6 – 0.75 = good and 0.75 – 1 = excellent (Cicchetti, 1994). In the HCP, when 5 minutes (post-censoring) of resting-state data were used to measure intrinsic connectivity defined within a common atlas of 264 regions (Power et al., 2011), the reliability was generally poor (mean ICC = .28, 95% CI [.28, .28]; 71% of edges were poor, 23% moderate, 6% good, and less than 1% excellent). As more resting-state data were added, the test-retest reliability continued to increase up to the limits of the dataset at 40 minutes (mean ICC = .54, 95% CI [.54, .54]; 26% of edges poor, 32% moderate, 25% good, and 18% excellent) (Figure 1A and left panel of Figure 2).

GFC, defined as exactly equal parts of task and resting-state data in the HCP, followed a similar pattern of increasing reliability with increasing data: 5 minutes of data exhibited poor reliability (mean ICC = .28, 95% CI [.28, .28]; 70% of edges were poor, 22% moderate, 7% good, and less than 1% excellent) but 40 minutes of data showed good reliability (mean ICC = .56, 95% CI [.55, .56]; 25% of edges were poor, 27% moderate, 24% good, and 23% excellent) (see Figures 1B, right panel of Figure 2). In addition, resting-state scans were not required to generate a reliable measure of GFC, as this pattern of increasing reliability held when task data alone were used to measure intrinsic connectivity (see supplemental Table S1). Furthermore, data from individual task scans alone generally resulted in poor reliability (see supplemental figures S2 and S3). A paired sample t-test comparing ICCs across all edges revealed that GFC is significantly more reliable than resting-state functional connectivity when both are estimated with 40 minutes of data (t(34715) = 140.01, p < .001). While significant, this difference is not likely meaningful as it represents only a 0.02 larger mean ICC in GFC. The low p-value is driven by the large number of degrees of freedom in the t-test comparing a mean ICC estimate from nearly 35,000 edges. Thus, we see the mean ICC of RSFC and GFC as statistically separable but practically equivalent.

This general improvement in reliability with increasing data was not unique to the HCP but was also observed in the population-representative Dunedin Study. The Dunedin Study resting-state scan of 8:16 minutes showed generally low reliability (mean ICC = .30, 95% CI [.30, .30]; 65% of edges were poor, 23% moderate, 10% good, and 2% excellent). The average reliability of the intrinsic connectivity estimates, however, improved by 49% when using GFC created in the Dunedin Study by combining the 8:16 minutes of resting-state data with 27 minutes (before censoring) of task data (mean ICC = .45, 95% CI [.45, .45]; 41% of edges were poor, 26% moderate, 19% good and 14% excellent) (Supplemental Figure S3).

### Influence of the global signal on reliability

GSR is still a controversial preprocessing step. Therefore, the main reliability analyses in the HCP were rerun without GSR to test the influence of GSR on ICCs. Mean ICCs for 5 minutes of data without GSR were poor for both RSFC (mean ICC = .31, 95% CI [.31, .31]; 68% of edges were poor, 27% moderate, 4% good, and < 1% excellent) and GFC (mean ICC = .35, 95% CI [.35, .35]; 60% of edges were poor, 31% moderate, 8% good, and 1% excellent) (see supplemental figure S1). In parallel fashion to the data processed with GSR, ICCs increased as scan length increased up to 40 minutes for both RSFC (mean ICC = .61, 95% CI [.61, .61]; 8% of edges were poor, 34% moderate, 41% good, and 16% excellent) and GFC (mean ICC = .63, 95% CI [.63, .63]; 8% of edges were poor, 30% moderate, 40% good, and 22% excellent). Despite the improvement in mean ICC with additional data with and without GSR, ICCs were significantly higher when GSR was not applied: 40 minutes of GFC had an average ICC that was 0.07 higher without GSR than when GSR was applied. The same pattern was present in RSFC: 40 minutes of resting-state without GSR was more reliable on average than with GSR (rest with GSR = .61, rest without GSR = .54). While higher reliabilities without GSR could be a sign of improved detection of true individual differences in intrinsic connectivity, they may also be a sign of a stable propensity to move within the scanner, creating correlated motion artifact in test and retest data (Power et al., 2016; van Dijk et al., 2012). Because the global signal is highly susceptible to motion artifact and there is no widely accepted method to isolate and remove just the artifactual component of the global signal while retaining the neural component, we adopted a conservative approach by applying GSR signal in all further analyses to ensure removal of motion related artifact present in the global signal (Power et al., 2018).

### Is there network specificity to improvements in reliability?

We next investigated network specificity in reliability by looking specifically at the reliability of edges within each functional network rather than the reliability of the entire correlation matrix in aggregate. As shown in Figure 3, there was clear network heterogeneity in reliability. While all networks improved in reliability from 10 minutes of data to 40 minutes, some networks improved more. With both RSFC and GFC, the limbic network had the lowest reliability at all scan lengths, improving from 10 minutes of data (RSFC: mean ICC = .22, 95% CI [.19, .25]; GFC: mean ICC = .30, 95% CI [.25, .34]) to 40 minutes of data (RSFC: mean ICC = .37, 95% CI [.33, .41]; GFC: mean ICC = .39, 95% CI [.35, .44]), but failing to escape a poor mean ICC. In contrast, networks that over-represent heteromodal association cortex (Margulies et al., 2016; Mesulam, 1998), namely the default mode network and fronto-parietal network, on average achieved moderate reliability with just 10 minutes of data for both RSFC (mean fronto-parietal network ICC = .53, 95% CI [.52, .53]; mean default mode network ICC = .57, 95% CI [.57, .58]) and GFC (mean fronto-parietal network ICC = .55, 95% CI [.54, .56]; mean default mode network ICC = .57, 95% CI [.57, .58]). Good reliability was achieved for these networks with 20 minutes of data for both RSFC (mean fronto-parietal network ICC = .62, 95% CI [.61, .62]; mean default mode network ICC = .66, 95% CI [.66, .67]) and GFC (mean fronto-parietal network ICC = .64, 95% CI [.63, .64]; mean default mode network ICC = .65, 95% CI [.65, .66]). With 40 minutes of data, these networks had mean reliabilities near the excellent range for both RSFC (mean fronto-parietal network ICC = .73, 95% CI [.72, .73]; mean default mode network ICC = .75, 95% CI [.75, .75]) and GFC (mean fronto-parietal network ICC = .74, 95% CI [.74, .75]; mean default mode network ICC = .77, 95% CI [.77, .77]). While there were some small differences in the network-specific reliabilities between RSFC and GFC (e.g., see dorsal attention network in Figure 3), network specific improvements with increasing scan length were similar for RSFC and GFC.

**Figure 3.**
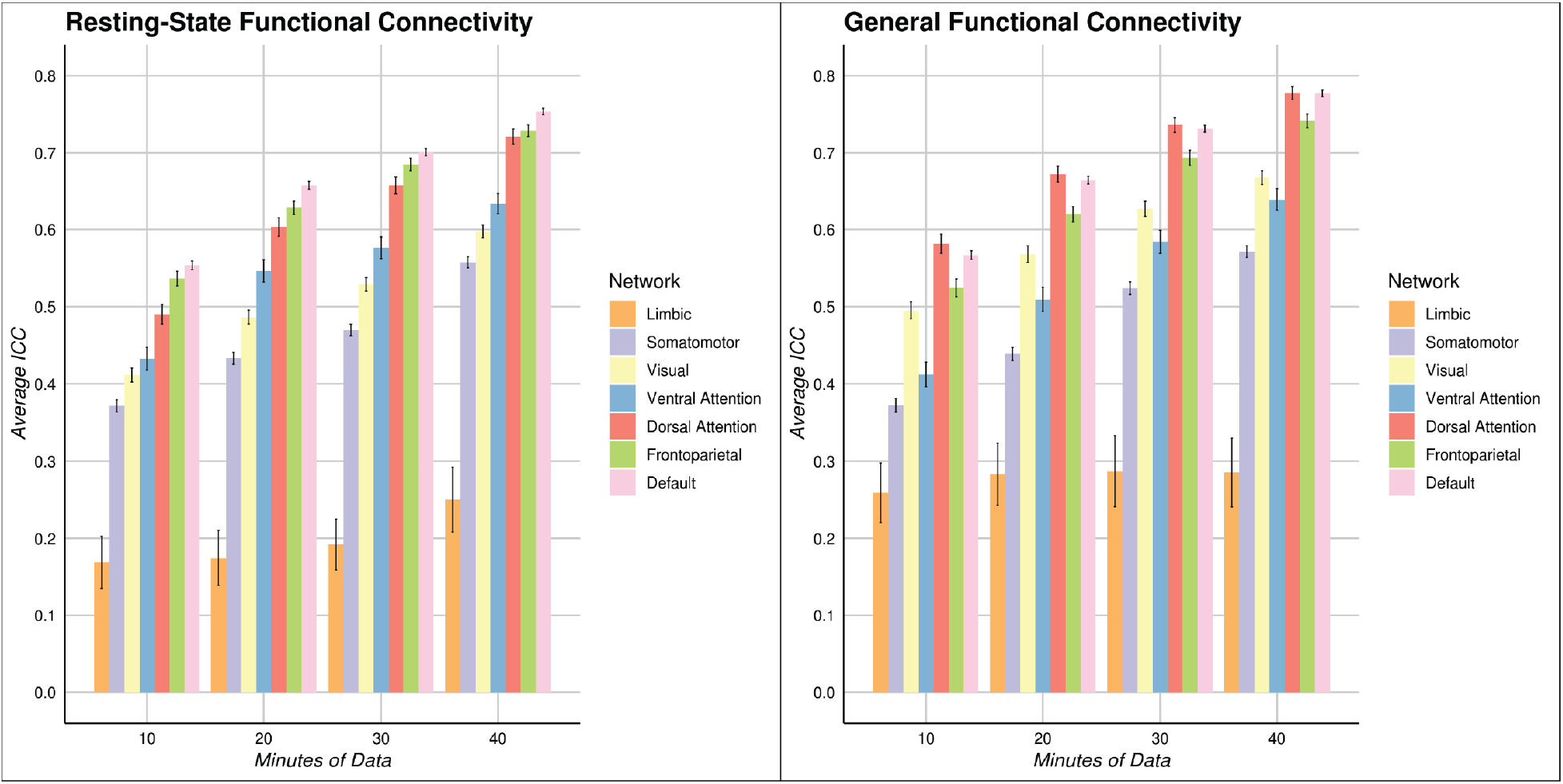
The improvement in test-retest reliability of intrinsic connectivity as a function of adding data varies across canonical functional networks. In the left panel mean ICC for RSFC are displayed for within-network connections across 7 previously defined networks (Yeo et al., 2011) for a variety of scan lengths. In the right panel the same data are displayed for GFC. There are clear and consistent differences between networks. The limbic network consistently has the lowest mean ICC, while the default mode network consistently has the highest mean ICC.

### What is the heritability of RSFC and GFC at different scan lengths?

As shown in Figure 4A, the heritability of intrinsic connectivity depended on scan length. The amount of variance for individual differences in RSFC attributable to the additive genetics (A) component of the ACE model increased consistently as the amount of data increased from 5 minutes (mean A component = .09, 95% CI [.09, .1]) to 40 minutes (mean A component = .22, 95% CI [.21, .23]). This represents an increase in the amount of variance in RSFC attributable to additive genetics of 138% as a function of increased scan length. This increase in variance explained by additive genetics was also present with GFC from 5 minutes (mean A component = .14, 95% CI [.13, .14]) to 40 minutes (mean A component = .28, 95% CI [.27, .29]). This represented an increase in the amount of variance in GFC attributable to additive genetics of 107% as a function of increased scan length. In addition, at equivalent scan lengths, GFC had significantly higher mean heritability across edges than RSFC (e.g., 5 minutes: t(945) = 10.11, p < .001) and 40 minutes: (t(945) = 14.75, p < .001)). Correspondingly, higher mean heritability in GFC than RSFC led to a greater percentage of heritable edges in GFC (see Figure 4B). In the ACE modeling, this pattern of increased heritability with increased scan time was associated with simultaneously decreased amount of variance explained by the E component, which comprises both non-shared environment and measurement error (Figure 4C). The E component decreased for both RSFC and GFC as data length increased from 5 minutes (RSFC mean E component = .81, 95% CI [.81, .82]; GFC mean E component = .77, 95% CI [.76, .77]) to 40 minutes (RSFC mean E component = .63, 95% CI [.63, .64]; GFC mean E component = .61, 95% CI [.60, .61]).

**Figure 4.**
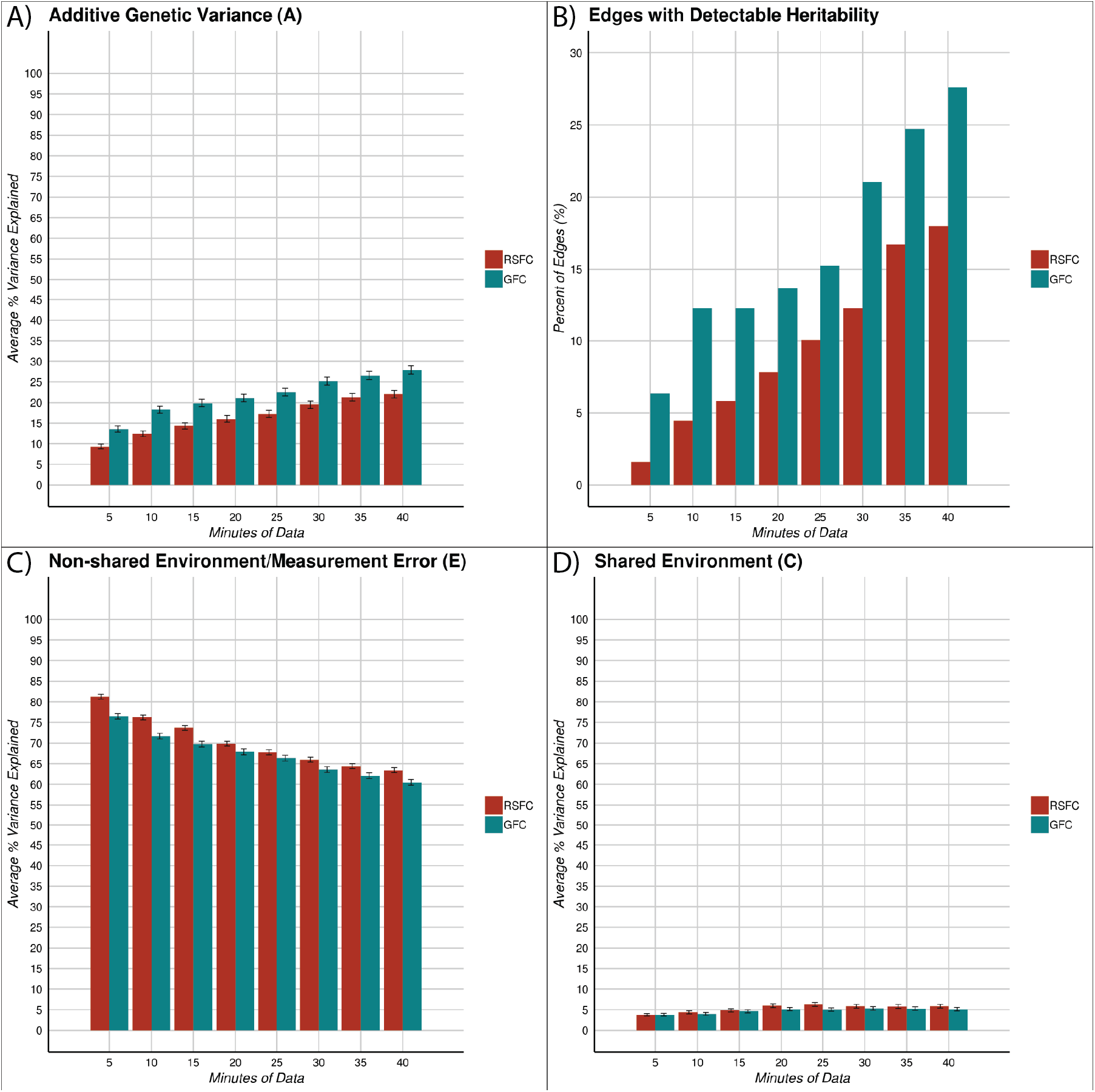
The heritability of RSFC and GFC vary as a function of scan length. Panels A, C and D display the mean A (additive genetics), E (non-shared environment + error) and C (shared environment) component estimates and 95% confidence intervals around those estimates, derived from ACE modeling of the twin data in the HCP. Scan length increases the A component and decreases the E component while having little effect on the C component. GFC also consistently has more variance attributable to the A component and lower variance attributable to the E component relative to RSFC. In panel B the % of heritable edges for RSFC and GFC are displayed across a variety of scan lengths. A heritable edge is defined as an edge with a lower bound of the 95% confidence interval that is larger than 0 in the ACE model.

While the A and E components of the ACE modeling changed substantially with scan length, the shared-environment (C) (Figure 4D) and twin-specific (T) (supplemental Figure S4) components were relatively stable with scan length. The variance attributable to the C and T components was similar for 5 minutes of RSFC (mean C component = .04, 95% CI [.03, .04]; mean T component = .06, 95% CI [.05, .06]) and 5 minutes of GFC (mean C component = .04, 95% CI [.03, .04]; mean T component = .06, 95% CI [.06, .07]) as well as with 40 minutes of RSFC (mean C component = .06, 95% CI [.05, .06]; mean T component = .09, 95% CI [.08, .09]) and 40 minutes of GFC (mean C component = .05, 95% CI [.05, .06]; mean T component = .06, 95% CI [.06, .07]).

### Exploring the connection between intrinsic connectivity and cognitive ability

CPM was used to predict cognitive ability from both RSFC and GFC in both the HCP and the Dunedin Study. In the first set of analyses, within-sample predictions were made using leave-one-out cross validation in the HCP and Dunedin Study datasets separately (Figure 5, left and center panels). In the HCP dataset, we found that individual differences in cognitive ability could be predicted from 40 minutes of RSFC (r^2^ = 4.2%, 95% CI [1.0%, 9.7%]) and from 40 minutes of GFC (r^2^ = 7.6%, 905% CI [2.7%, 14.2%]) (see Figure 5 and supplemental Table S3). These results were replicated and extended in the Dunedin Study using scan lengths that are more representative of existing datasets than the HCP. Cognitive ability could be predicted from the 8-minute RSFC data (r^2^ = 6.8%, 95% CI [3.5%, 11.3%]) as well as from 33 minutes of GFC (r^2^ = 10.2%, 95% CI [5.9%, 15.0%]; all available data; supplemental Table S4). Although in comparison with RSFC, GFC had 81% greater predictive utility in the HCP dataset and 50% greater predictive utility in the Dunedin Study dataset, the statistical comparison of the correlations was not statistically significant in either the HCP (GFC r^2^ = 7.6%, RSFC r^2^ = 4.2%, Steiger’s z = 1.317, p = .188) or the Dunedin Study (GFC r^2^ = 10.2%, RSFC r^2^ = 6.8%, Steiger’s z = 1.72, p = 0.085). In addition, scan length had little effect on within-sample predictive utility (see supplemental figure S5).

**Figure 5.**
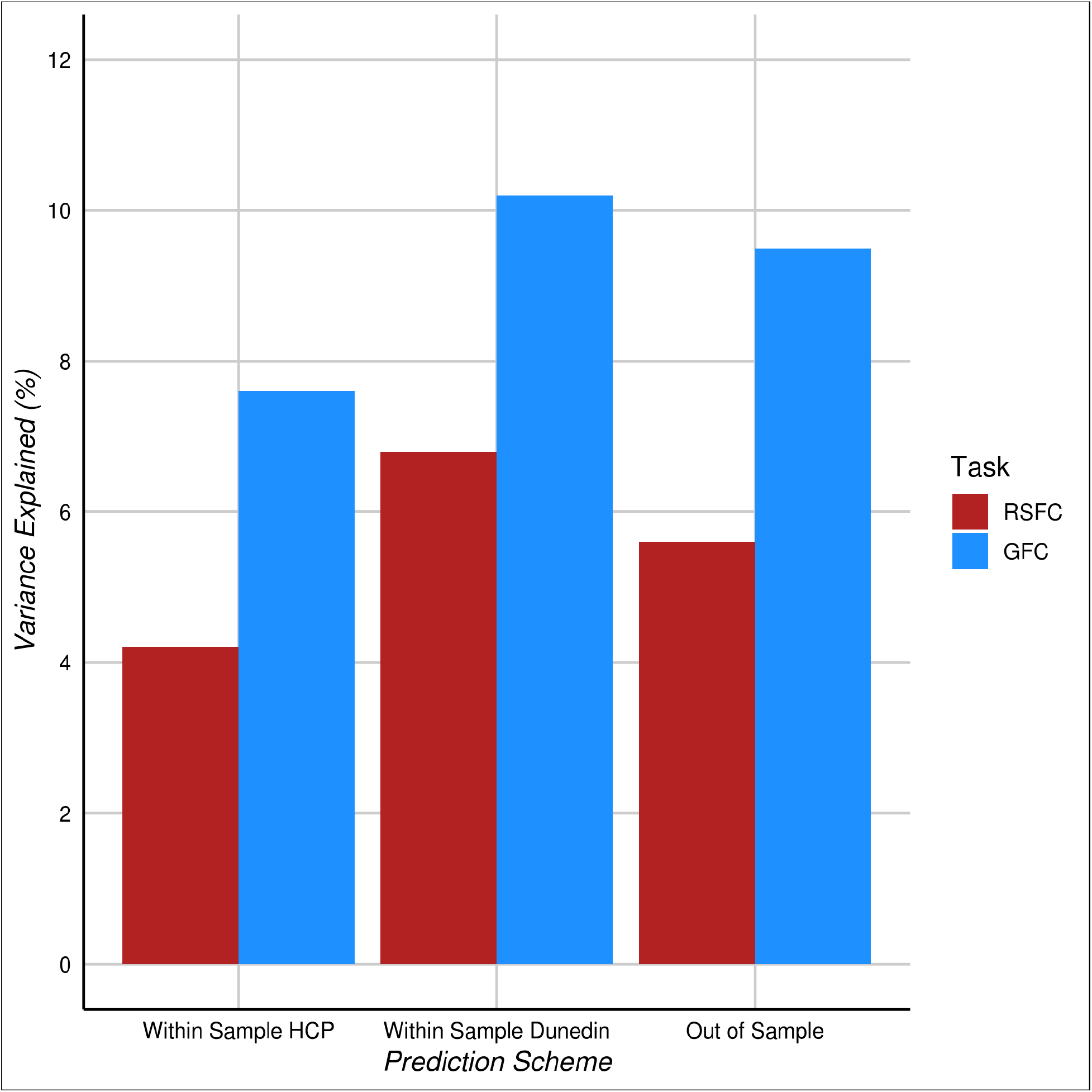
GFC is better than RSFC at predicting cognitive ability both within and between samples. Results from CPM models predicting cognitive ability from RSFC and GFC. The x-axis displays predictions from leave-one-out cross validation within sample and out-of-sample models trained using the Dunedin Study dataset and tested using the HCP dataset. Predictive utility is displayed as % variance explained (r^2^).

While leave-one-out cross validation has been widely used with CPM (Finn et al., 2015; Noble et al., 2017), the gold standard test of a model’s generalizable predictive utility is its ability to predict a phenotype in an independent sample (Bzdok and Yeo, 2017; Whelan and Garavan, 2014; Yarkoni and Westfall, 2017). This is particularly important for GFC, as it purports to measure a generalizable feature of intrinsic connectivity and thus correlations between individual differences in GFC and phenotypes of interest should be shared across independent samples with different batteries of fMRI tasks. Therefore, we used CPM to train two models (one with rest and one with GFC) to predict cognitive ability in the larger Dunedin Study dataset and tested generalizability by measuring the ability of these models to predict cognitive ability in the HCP dataset. The models built with RSFC (r^2^ = 5.6%, 95% CI [1.8%, 11.9%], p, q < .001) and GFC (r^2^ = 9.5%, 95% CI [4.0%, 16.3%], p, q < .001) from the Dunedin Study dataset both successfully predicted cognitive ability in the HCP dataset (Figure 5 right panel and Figure 6 left panel). However, GFC had greater out-of-sample predictive utility than RSFC (GFC r^2^ = 9.5%, RSFC r^2^ = 5.6%, Steiger’s z = 2.57, p = 0.010). This provides evidence that GFC is a more generalizable measure across independent samples than RSFC.

**Figure 6.**
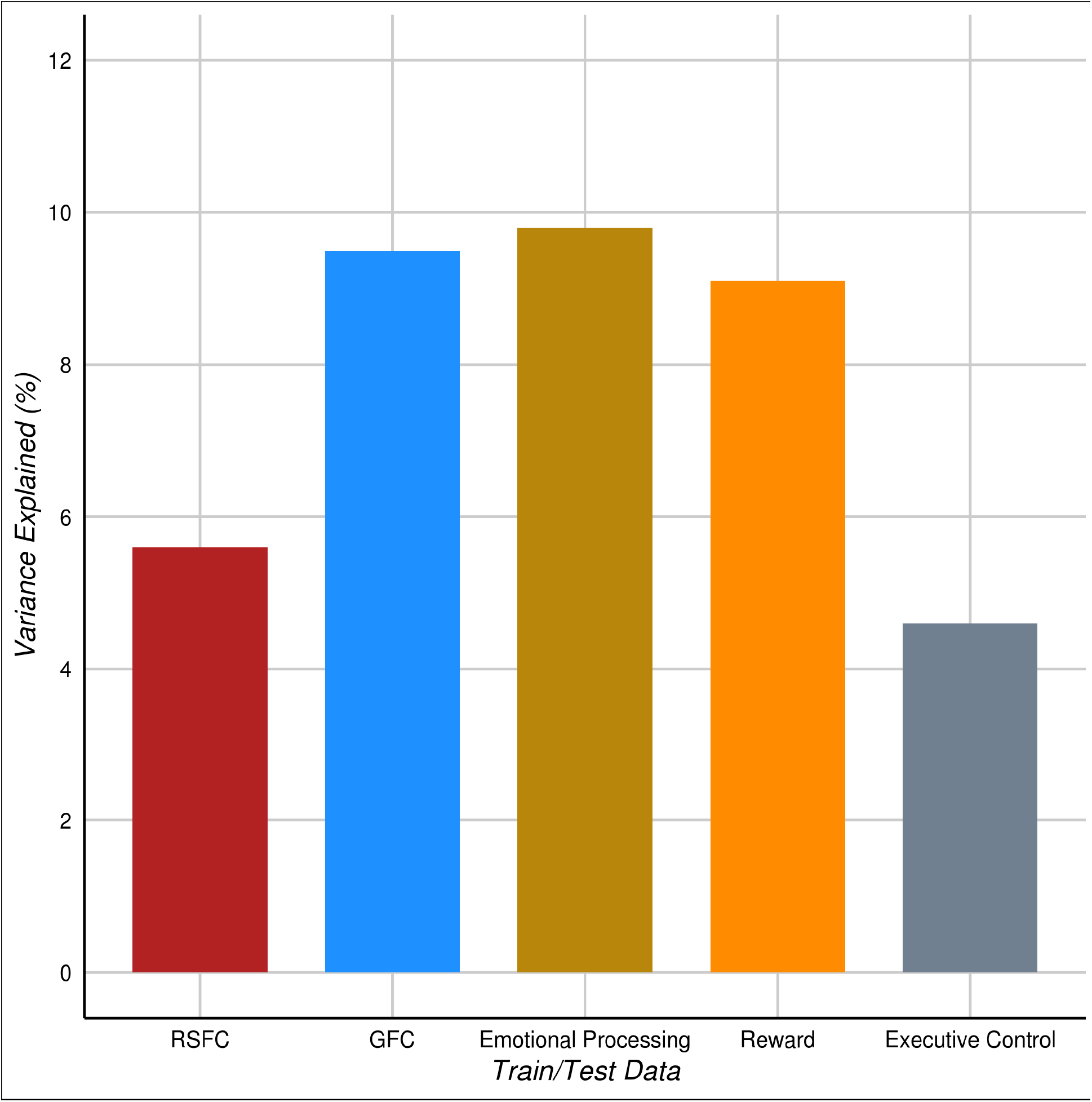
Out-of-sample prediction of cognitive ability for GFC is better than RSFC and as good as task-derived intrinsic connectivity. All models were trained on intrinsic connectivity data from the Dunedin Study and tested using data from the HCP. Models were trained and tested with the same type of data. With task data this meant that models were trained and tested with tasks that have a comparable (parallel) task in both the HCP and Dunedin study. Predictive utility is displayed as % variance explained (r^2^).

### Does GFC predict cognitive ability better than intrinsic connectivity measured from specific tasks?

Recent research has suggested that intrinsic connectivity estimates derived from task data are better at predicting cognitive ability than estimates derived from resting-state data (Greene et al., 2018). Our above finding that GFC outperforms RSFC at predicting cognitive ability is in line with this observation. However, given that many researchers collect different tasks and resting-state scans with different “task” demands (e.g., eyes-closed, fixation, etc.), we next investigated if GFC could perform as well or better than specific tasks at out-of-sample prediction of cognitive ability. We constructed five prediction models that were trained to predict cognitive ability using the Dunedin Study dataset (the population-representative sample with the gold-standard WAIS IQ test) and tested for generalizability using the HCP dataset (see supplemental figure S6 for the converse, models trained in HCP and tested in the Dunedin Study). Two models were carried over from the previous analyses (RSFC and GFC) and three additional task-specific models were added. These consisted of models trained and tested on three parallel tasks available in both the Dunedin Study and HCP (see methods for details). All models trained using the Dunedin Study dataset were able to significantly predict cognitive ability in the HCP dataset (all FDR q < .05; Figure 6 and supplemental Table S5). Prediction of cognitive ability from RSFC data (r^2^ = 5.6%, 95% CI [1.8%, 11.9%], p, q < .001), GFC data (r^2^ = 9.5%, 95% CI [4.0%, 16.3%], p, q < .001), emotional processing data (r^2^ = 9.8%, 95% CI [4.3%, 17.1%], p < .001, q = .001), reward processing data (r^2^ = 9.1%, 95% CI [3.6%, 16.5%], p < .001, q = .001) and executive function data (r^2^ = 4.6%, 95% CI [1.0%, 10.8%], p < .001, q = .001) all successfully generalized from the Dunedin Study dataset to the HCP dataset.

Comparing the relative performance of models trained with each type of data revealed that GFC was generalizable and performed as well or better at out-of-sample prediction of cognitive ability compared to all tasks (Figure 6). Predictions of cognitive ability from GFC performed better than RSFC (GFC r^2^ = 9.5%, RSFC r^2^ = 5.6%, Steiger’s z = 2.57, p = 0.01) and executive function data (GFC r^2^ = 9.5%, executive function r^2^ = 4.6%, Steiger’s z = 2.19, p = .03), and performed as well as emotional processing data (GFC r^2^ = 9.5%, emotion processing r^2^ = 9.8%, Steiger’s z = -.10, p = .92) and reward data (GFC r^2^ = 9.5%, reward r^2^ = 10.2%, Steiger’s z = -.31, p = .76). Out-ofsample prediction was largely unaffected by the amount of data used in constructing the intrinsic connectivity matrices but was adversely affected when task regression was removed (see supplemental Table S2).

## Discussion

Here we present GFC as a method for readily improving the reliability of estimating individual differences in intrinsic connectivity by combining resting-state and task data. Across two independent datasets we found that scan length has a significant impact on the reliability of intrinsic connectivity estimates regardless of whether the data came solely from resting-state scans (i.e., RSFC) or from a combination of resting-state and task scans (i.e., GFC). GFC, the addition of task scans to short resting-state scans, substantially increased the reliability of intrinsic connectivity over and above short resting-state scans alone. Moreover, we found that the improved reliability from aggregating data resulted in higher estimates of the heritable variation in intrinsic connectivity, and that this gain was larger for GFC than RSFC. In addition, GFC consistently performed better than RSFC and performed as well or better than intrinsic connectivity from individual task scans at predicting individual differences in a complex human trait, namely cognitive ability.

These findings have several implications for current and future neuroimaging research because they demonstrate that 1) reliable and heritable measurement of individual differences in the functional architecture of the brain is not only achievable in niche, specialty datasets with hours of resting-state scans, but also in many existing datasets if GFC is adopted; 2) mapping individual differences in behavior and cognition may be improved by adopting GFC, as demonstrated by generalizable predictive utility when studying cognitive ability; and 3) to the extent that commensurate gains in reliability and heritability may be achieved in other datasets, our results provide a reference that researchers can use to roughly estimate the gain in reliability they may achieve by adopting GFC (Figures 1, 2, S2 and S3). We next elaborate on each of these points.

### Data aggregation across task and resting-state scans boosts reliability

We found that GFC, derived from the combination of resting-state and task scans, can be a reliable measure of individual differences in intrinsic connectivity. Consistent with previous studies (Birn et al., 2013; Finn et al., 2015; Laumann et al., 2015), we showed that the test-retest reliability of intrinsic connectivity is highly dependent on the amount of data collected. This is true of RSFC as well as GFC. With only 5-10 minutes of data, both RSFC and GFC show poor reliability in the HCP and Dunedin Study datasets. Only with 30-40 minutes of data do RSFC and GFC begin to broadly display good reliability. These findings follow directly from past studies that have shown reliability depends on scan length (Anderson et al., 2011; Birn et al., 2013; Hacker et al., 2013), that task and resting-state data share a large proportion of variance (Cole et al., 2014; Geerligs et al., 2015), and that both task and resting-state data measure common individual differences in intrinsic connectivity (Gratton et al., 2018). Some functional networks, however, can achieve good reliability with relatively little data. For example, the default mode and fronto-parietal networks achieved good reliability for within-network connections with just 20 minutes of RSFC or GFC (see figure 3). In contrast, the limbic network failed to achieve even fair levels of reliability with up to 40 minutes of data. These findings suggest that the amount of data needed to derive reliable estimates of intrinsic connectivity for individual differences research will vary depending on the network of interest.

Importantly, the current reality is that most neuroimaging studies have only 5-10 minutes of resting-state data, which are further limited by sampling variability, motion-artifacts, and other sources of noise (Gordon et al., 2017; Gratton et al., 2018; Power et al., 2014). Given the reliability estimates reported here, it is likely that in many studies, individual differences in intrinsic connectivity will be unmeasurable with resting-state data alone (Anderson et al., 2011; Hacker et al., 2013). Fortunately, it is common practice for researchers to collect 10-40 minutes of task fMRI in addition to resting-state. Therefore, if researchers adopt a broader definition of intrinsic connectivity, such as GFC, they can combine task and resting-state data to achieve a reliable measure of individual differences in intrinsic connectivity.

### Data aggregation across task and resting-state scans boosts heritability

We found that individual differences in intrinsic connectivity in both RSFC and GFC were significantly attributable to additive genetic influences. With 40 minutes of data, 22% of the variance in RSFC and 28% of the variance in GFC was attributable to additive genetic effects. With equivalent amounts of data, GFC produced higher heritability estimates than RSFC, and the influence of additive genetic effects was detectable in a larger proportion of functional connections (Figure 4). In addition, we found that scan length had an impact on the heritability estimates of intrinsic connectivity in both RSFC and GFC. The average amount of variance in intrinsic connectivity that was attributable to additive genetic effects more than doubled in both RSFC (increase of 138%) and GFC (increase of 107%) as scan length increased from 5 to 40 minutes. While the heritability of intrinsic connectivity has been investigated before (Adhikari et al., 2018; Ge et al., 2017), the link between scan length and heritability has not been described. Although a reliable measure does not have to be heritable, a measure cannot have high heritability without high reliability (Ge et al., 2017). That is, measurement reliability places a ceiling on the heritability estimate of a phenotype. Therefore, our finding that the heritability of intrinsic connectivity increases with increasing reliability suggests that the low reliability of intrinsic connectivity in short fMRI scans limits estimations of the true underlying heritability of intrinsic connectivity in many datasets. More specifically, our finding suggests that researchers should consider the amount of fMRI data available for estimation of intrinsic connectivity when planning and evaluating imaging genetics research.

Our results have additional implications for genetically-informed fMRI research. Neuroimaging measures have been considered promising intermediate phenotypes that would bring researchers closer to the mechanisms through which genetics lead to heritable psychological traits (Braskie et al., 2011; Hariri and Weinberger, 2003; Hasler and Northoff, 2011; Meyer-Lindenberg and Weinberger, 2006). To uncover these links, large datasets like ENIGMA and the UK Biobank (Sudlow et al., 2015; Thompson et al., 2014), have been used to conduct genome wide association studies (GWAS) of MRI-based phenotypes. While these investigations have had success in finding genetic variants linked to structural MRI measures (Adams et al., 2016; Hibar et al., 2017, 2015; Stein et al., 2012), they have largely failed to find significant genetic correlates of functional intrinsic connectivity measures (Elliott et al., 2017). One reason for this failure may be the low reliability and heritability of the short resting-state scans typically used to derive individual differences measures of RSFC in these studies. Our results suggest that future GWAS may be better powered to find genetic variants linked to the functional architecture of the brain by adopting GFC rather than RSFC from short resting-state scans.

### How general is general functional connectivity?

Previous research has shown that the predictive utility of intrinsic connectivity can be higher when derived from task data than when derived from resting-state data (Greene et al., 2018). Some task states in particular may reveal individual differences in intrinsic connectivity that aid in predictive utility (Finn et al., 2017). While these findings provide insight into individual differences in intrinsic connectivity and a new use for task data, they pose a problem for replication and cumulative aggregation of findings across datasets. While most fMRI studies collect task data, very few collect the same tasks. Our findings here suggest that GFC may provide a solution to this problem. First, we found that like task-derived intrinsic connectivity, GFC can outperform RSFC when predicting within-sample and out-of-sample variability in cognitive ability (Figures 5 and 6). Second, we found that GFC performed as well or better than specific tasks alone at predicting cognitive ability out-of-sample. Critically, in these comparisons GFC was constructed with a combination of different tasks in the training and testing datasets suggesting that GFC may measure a common, generalizable backbone of intrinsic connectivity that can be applied and aggregated across independent datasets with different task batteries (supplemental Tables S6 and S7). While further research is needed to replicate these findings, they nevertheless suggest that GFC may provide a generalizable, practical framework that researchers with resting-state and task data can use to drive a cumulative, predictive neuroscience of individual differences (Button et al., 2013; Szucs and Ioannidis, 2017).

### Is general functional connectivity too heterogeneous?

An understandable reluctance in implementing GFC could be that it would introduce heterogeneity in the estimation of intrinsic connectivity. Whereas RSFC is thought to be a common framework with generalizability across datasets, the combination of different sets of tasks in different datasets may introduce additional variability. Below, we provide three reasons why we think this is less problematic than it first appears. We further provide specific recommendations and guidelines for adopting and estimating GFC and point to features of our analyses that support the validity of GFC as a robust and reliable measure of individual differences in intrinsic connectivity.

First, resting-state data are by nature heterogenous, and the resting-state is its own type of task (Buckner et al., 2013). Researchers collect resting-state data under different conditions in which participants close their eyes, have their eyes open, or fixate. These differences in resting-state conditions influence intrinsic connectivity (Patriat et al., 2013). Despite these differences, all approaches are collectively referred to as resting-state scans. In addition, factors like thought content (Christoff et al., 2009; Hurlburt et al., 2015), caffeine intake (Wong et al., 2012), recent cognitive tasks (Waites et al., 2005), and falling asleep (Deco et al., 2014) can introduce further heterogeneity that can bias intrinsic connectivity estimates. It has even been estimated that approximately 30% of participants fall asleep within the first three minutes of a resting-state scan (Tagliazucchi and Laufs, 2014). Nevertheless, resting-state scans measure a common set of functional networks (Yeo et al., 2011), and display trait-like features driven by stable factors such as genetics (Glahn et al., 2010) and structural connectivity (Honey et al., 2009). For these reasons resting-state data represent substantial promise as an individual-differences measure if enough data are collected to average out sampling variability (Gratton et al., 2018). But, as already stated, few researchers collect enough data to fulfill this promise. Connectivity estimates from task data are also shaped by many of these stable factors (Cole et al., 2014; Krienen et al., 2014), and data from different tasks predominantly measure overlapping individual differences that are only weakly influenced by task demands (Gratton et al., 2018). Admittedly, GFC will vary, to some extent, across samples because each study collects a unique combination of resting-state and task data. However, RSFC will vary too. Moreover, we found that disparate task and resting-state scans collectively measure reliable, heritable individual differences with generalizable out-of-sample predictive utility in the form of GFC.

Second, the benefits of GFC converged across two different samples, with different scanners, scanning parameters, and demographics. The HCP represents a healthy sample of highly-educated individuals (Van Essen et al., 2013), while the Dunedin Study represents a population-representative birth cohort with a broad range of mental and physical health conditions, socioeconomic status, and full representation of variability in many complex traits (Poulton et al., 2015). Across samples, the data came from 11 different tasks (HCP = 7, Dunedin Study = 4), many with unrelated cognitive demands, different scan lengths, stimuli, behavioral requirements, and instructions. The data were processed with different preprocessing schemes and software (“minimally preprocessed” in the HCP and custom scripts in the Dunedin Study). While the amount of scan time was tightly controlled in the HCP analyses (i.e., equal time after censoring), in the Dunedin Study motion censoring led to unequal scan lengths across participants as is the case in many analyses of intrinsic connectivity. Despite these many differences between datasets, the results demonstrate that GFC can achieve good test-retest reliability, inter-rater reliability (different preprocessing schemes), out-of-sample reliability (convergence across datasets and out-of-sample prediction), and parallel forms reliability (different tasks in each sample) (Dubois and Adolphs, 2016). While we present evidence in favor of the generalizability of GFC, future research should further investigate the heterogeneity in GFC to find combinations of tasks that most efficiently estimate individual differences in intrinsic connectivity (Finn et al., 2017; Shah et al., 2016).

Third, reliability fundamentally limits the ability to detect associations between any two measures (Nunnally Jr., 1970; Vul et al., 2009). Therefore, any investigation mapping intrinsic connectivity to individual differences in behavior, cognition, or disease is fundamentally limited by the reliability of the intrinsic connectivity measures. Relatedly, statistical power to detect true effects depends on the anticipated effect size, which is in turn dependent on reliability of each measurement (Kanyongo et al., 2007; Williams and Zimmerman, 1989). Given that many studies only have 5-10 minutes of resting-state data and consequently poor reliability of intrinsic connectivity measures, they have drastically reduced statistical power (Button et al., 2013; Ioannidis, 2008, 2005), and likely tenuous correlates of individual differences unless they adopt a framework like GFC. Previous research has found that collectively a large number of unreliable edges can achieve multivariate reliability that results in predictive utility (Noble et al., 2017). While multivariate predictive utility may be achievable in studies with poor univariate edge-wise reliability (Figure S5), the interpretability and clinical utility of these studies will be diminished by low univariate reliability. This is because interpretability and clinical utility depend on isolating specific brain areas or functional connections that can be said to be “most” important to the phenotype of interest. This cannot be done without decent univariate reliability because accurate estimation of true parameters in a prediction model depends on reliability (Cremers et al., 2017; Vul et al., 2009; Yarkoni and Westfall, 2017). For example, if researchers want to characterize biomarkers of mental illness that could be targeted with interventions like transcranial magnetic stimulation (Fox et al., 2014) or real-time neurofeedback (Caria et al., 2012), they will need to isolate specific circuits that with confidence can be said to be “altered” and relevant to the disease process in mental illness. Actionable identification of these biomarkers will only be possible with adequate univariate reliabilities. GFC provides a practical solution to this problem, while also addressing power and reliability issues that have been identified more generally in neuroscience (Button et al., 2013; Szucs and Ioannidis, 2017) and beyond (Aarts et al., 2015; Errington et al., 2014; Ioannidis, 2005; Ioannidis et al., 2001; Munafò et al., 2017). GFC provides a tool for researchers to repurpose existing and future datasets to better contribute to a cumulative science of individual differences (Dubois and Adolphs, 2016) with clinical relevance (Fox, 2010). For these reasons, we think that in many studies the gains in reliability offered by GFC will outweigh potential task-induced “heterogeneity.”

### Recommendations

Based on the above summary, we generally recommend that researchers seeking to maximize the reliability and generalizable predictive utility of GFC should aggregate as much data as possible, across all rest and task data, that they have at their disposal. Our results further suggest that with a minimum of 20 minutes of GFC data, researchers can achieve good reliability in some networks of broad interest (e.g., default, frontoparietal and dorsal attention networks). Furthermore, a strict balance of the amount of data from each task and resting-state scan is not necessary to achieve the benefits of GFC. This is demonstrated by the increased reliability of GFC despite unequal task lengths in the Dunedin Study dataset and in the HCP dataset by continued improvement in reliability despite timepoints being selected at random from the remaining data after reaching a minimum of 25 minutes of data. However, we urge caution in applying GFC to datasets that are heavily skewed (> 50%) towards resting-state or one specific task. Although we expect that deriving GFC from this type of dataset will boost reliability, the generalizability of these findings to GFC derived from other datasets may be limited because of the overrepresentation of a specific type of data. Lastly, our results suggest that researchers developing new protocols may benefit from collecting multiple short task scans rather than a single longer task scan (supplemental tables S1 and S5).

### Conclusion

Here we propose GFC as a method for improving the reliability of estimating individual differences in the intrinsic architecture of functional networks. We demonstrate that, when the amount of data available for analysis is held constant, the combination of resting-state and task data is at least as reliable as resting-state alone. Additionally, when the same amount of data is available, GFC produces higher heritability estimates than RSFC. Many researchers who have less than 25 minutes of resting-state data but have additionally collected task data on the same participants can immediately boost reliability, heritability, and power by adopting GFC as a measure of intrinsic connectivity when studying individual differences. Our findings also suggest that future data collection should consider naturalistic fMRI (Hasson et al., 2010, 2004; Huth et al., 2016; Lahnakoski et al., 2012; Vanderwal et al., 2017) and engaging tasks in addition to resting-state scans when generating data for estimating intrinsic connectivity. This may be especially true when studying children, elderly, or mentally ill participants, as these groups often cannot lie still for 25 minutes or more of resting-state scanning (Power et al., 2012; Satterthwaite et al., 2013; Yuan et al., 2009). Our current findings also suggest that researchers need not think of collecting resting-state and task data as a zero-sum competition for scan time when designing studies. Instead, our findings demonstrate that task and resting-state data provide complementary measures that together can lead to more reliable, heritable measurement of individual differences in intrinsic connectivity. We think that collecting adequate amounts (i.e., > 25 minutes) of high-quality fMRI data should be emphasized over the traditional task-rest dichotomy, when individual differences are of interest. If fMRI is going to be a part of a cumulative neuroscience of individual differences with clinical relevance, psychometric measurement quality, especially reliability, must be at the forefront of study design.

## Acknowledgement

The Dunedin Multidisciplinary Health and Development Study is supported by the NZ HRC and NZ MBIE. This research was supported by National Institute on Aging grant R01AG032282, R01AG049789, and UK Medical Research Council grant MR/P005918/1. Additional support was provided by the Jacobs Foundation. MLE is supported by the National Science Foundation Graduate Research Fellowship under Grant No. NSF DGE-1644868. Thank you to members of the Advisory Board for the Dunedin Neuroimaging Study. We thank the Dunedin Study members, unit research staff, and Study founder Phil Silva. The study protocol was approved by the institutional ethical review boards of the participating universities. Study members gave informed consent before participating. The authors declare no competing financial interests.

## Author Contributions

Conceptualization, M.L.E.; Methodology, Software and Formal Analysis, M.L.E., A.R.K and M.C.; Data Curation, M.L.E., A.R.K., A.C., and T.E.M.; Resources, T.R.M., R.K., S.R., R.P., A.C., T.E.M. and A.R.H., Writing, M.L.E., A.R.K., M.J.K., M.C., T.R.M., R.K., D.I., S.R., R.P., A.C., T.E.M. and A.R.H.; Supervision, A.C., T.E.M. and A.R.H.; Funding Acquisition, R.P., A.C., T.E.M. and A.R.H.

