## Supplemental Information for "General Functional Connectivity: shared features of resting-state and task fMRI drive reliable and heritable individual differences in functional brain networks"

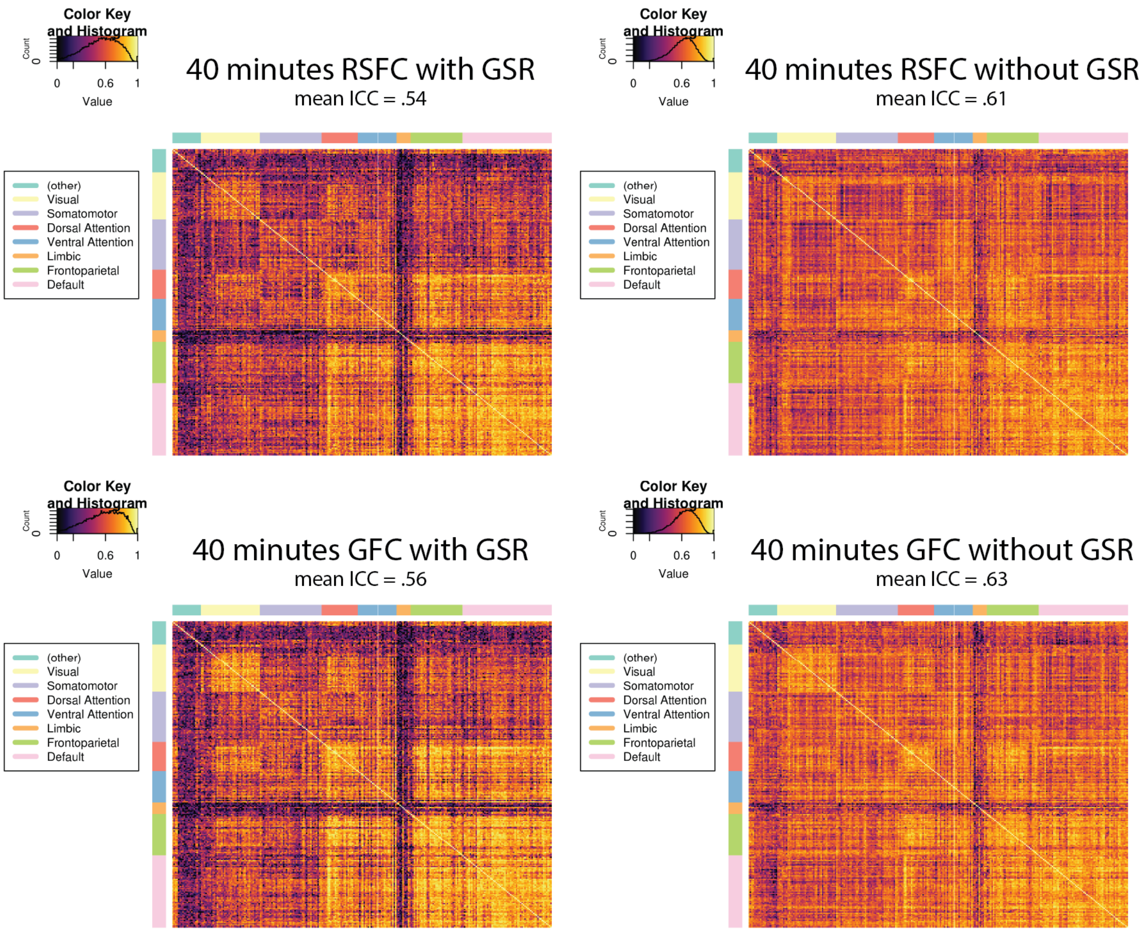


**Figure S1.** Influence of GSR on ICC matrices for RSFC (top panel) and GFC (bottom panel). The matrices in the left panel are constructed from data that has been processed with global signal regression (GSR), and those in the right panel when there has been no GSR. There is clearly higher reliability on average when GSR in not applied. However, because the global signal has been associated with artifactual changes in functional connectivity all primary analyses reported in the main text include GSR.

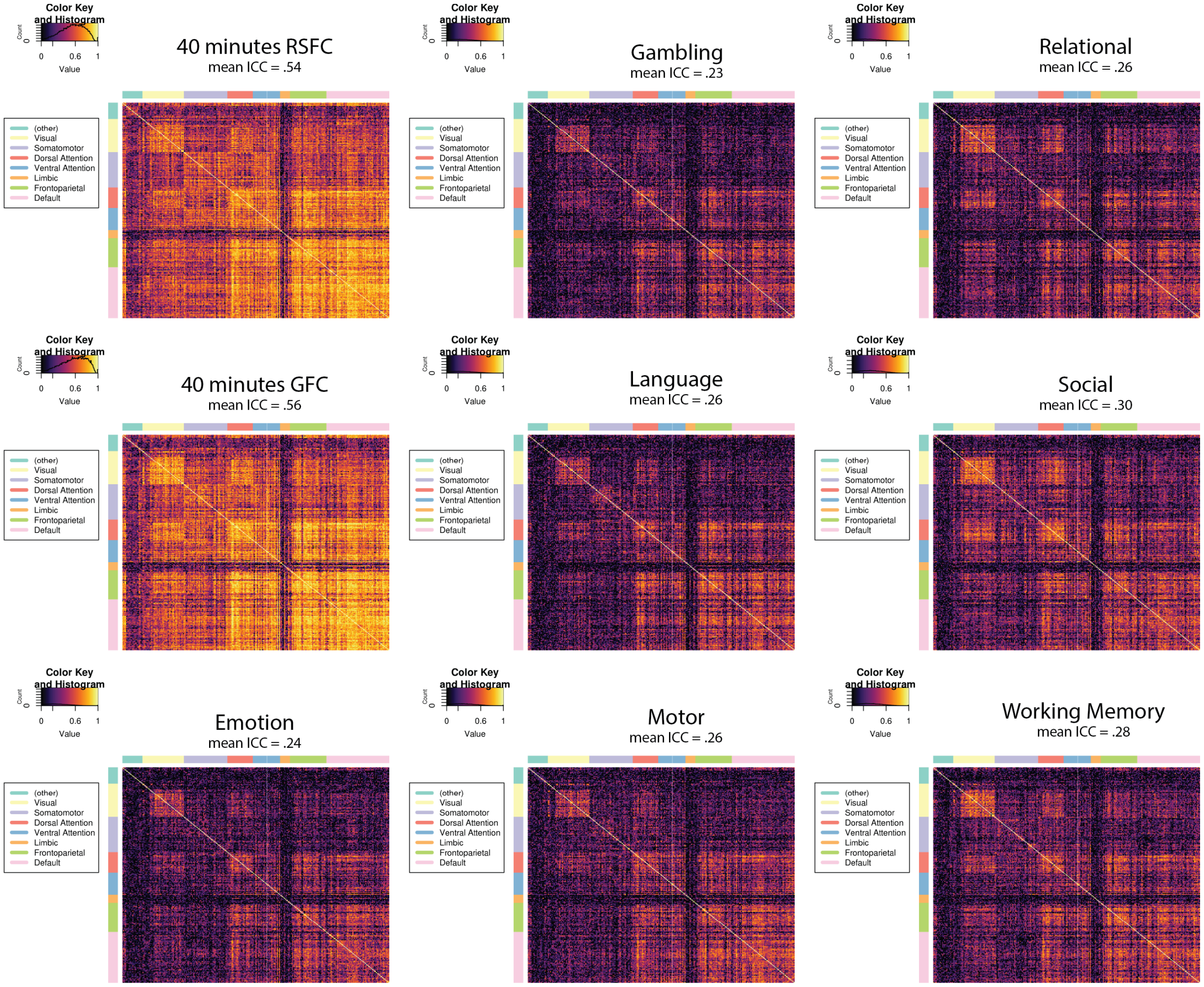


**Figure S2.** ICC matrices for RSFC, GFC, and task-specific intrinsic connectivity estimates from the HCP dataset. This figure demonstrates that increased GFC reliability does not rely on a particular task, as intrinsic connectivity from all tasks when investigated in isolation have poor reliabilities. Only when tasks are combined into longer scans to estimate GFC do they achieve better reliabilities. This suggests that tasks share common individual differences in intrinsic connectivity that can be more reliably measured when relatively short task scans are combined than when they are investigated in isolation.

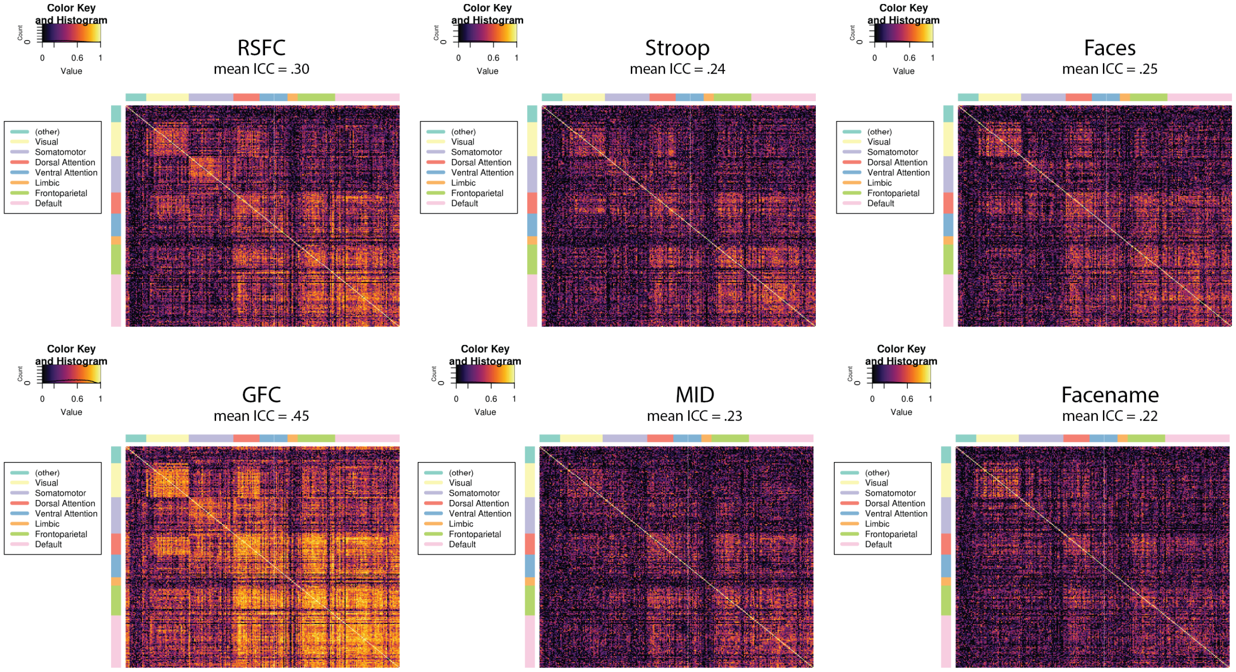


**Figure S3.** ICC matrices for RSFC, GFC, and task-specific intrinsic connectivity estimates from the Dunedin Study dataset, which effectively replicate the patterns displayed in Figure S2.

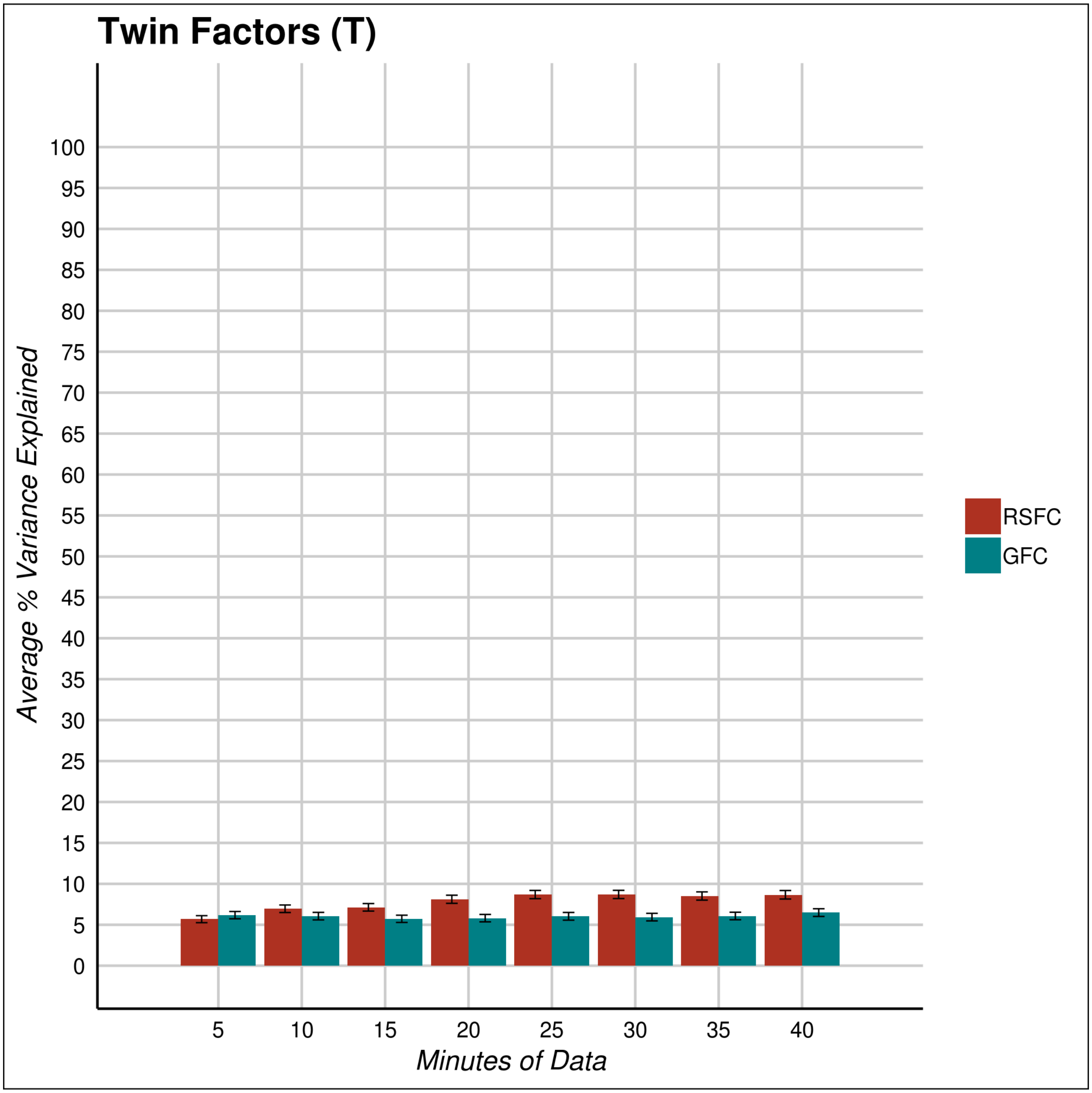


**Figure S4.** The percentage of variance explained by twin specific factors as a function of scan length from the ACE modeling. Scan length has little effect on the twin specific estimates.

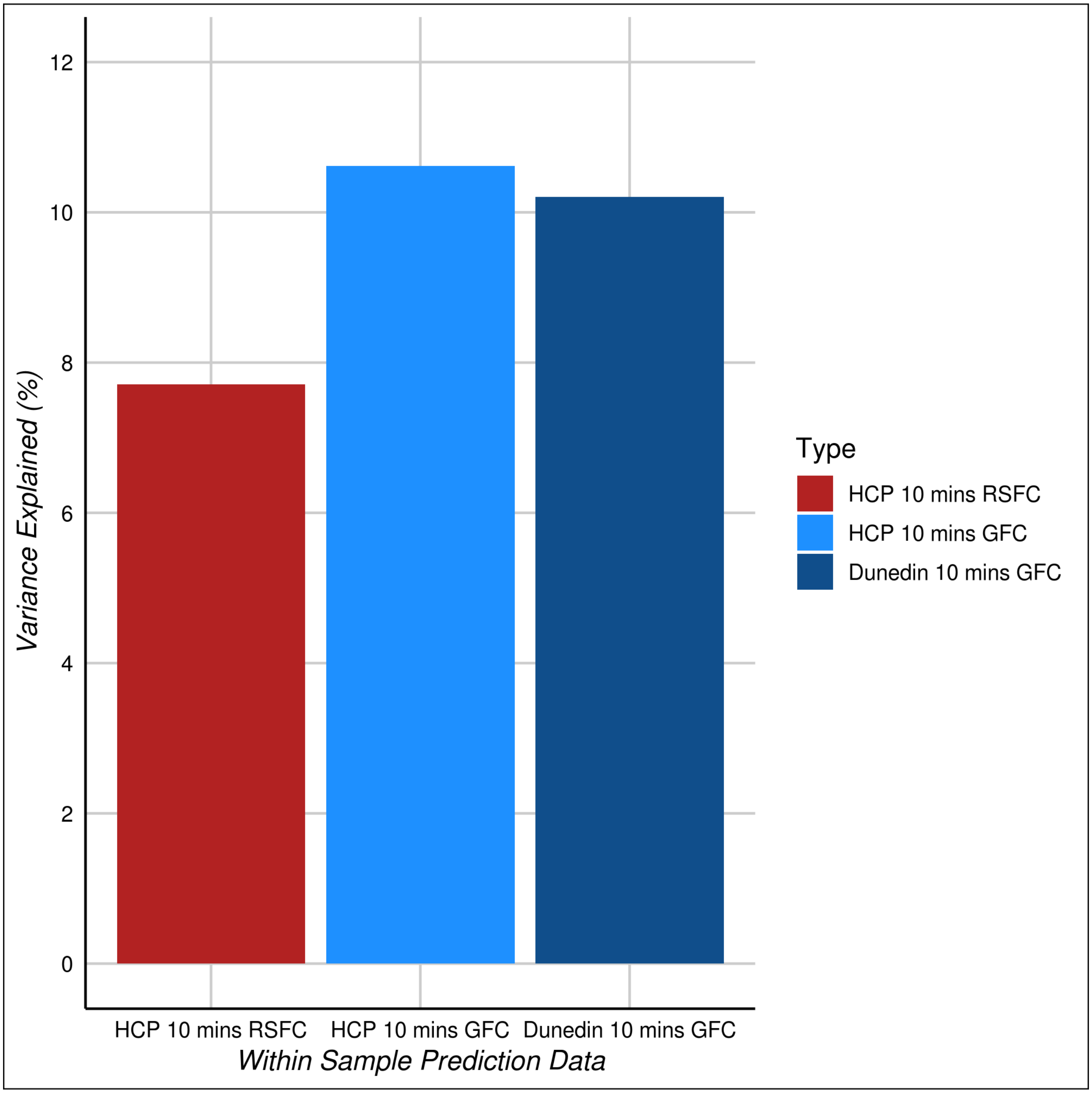


**Figure S5.** Bar plot displaying the predictive utility of RSFC and GFC with only 10 minutes of data. Predictions are all made within-sample using. Model performance is comparable to within-sample performance when using 2-3 times as much data (Figure 5). This indicates the modest impact increasing scan length has on predictive utility despite its large effect on reliability (Figure 2) and heritability (Figure 4).

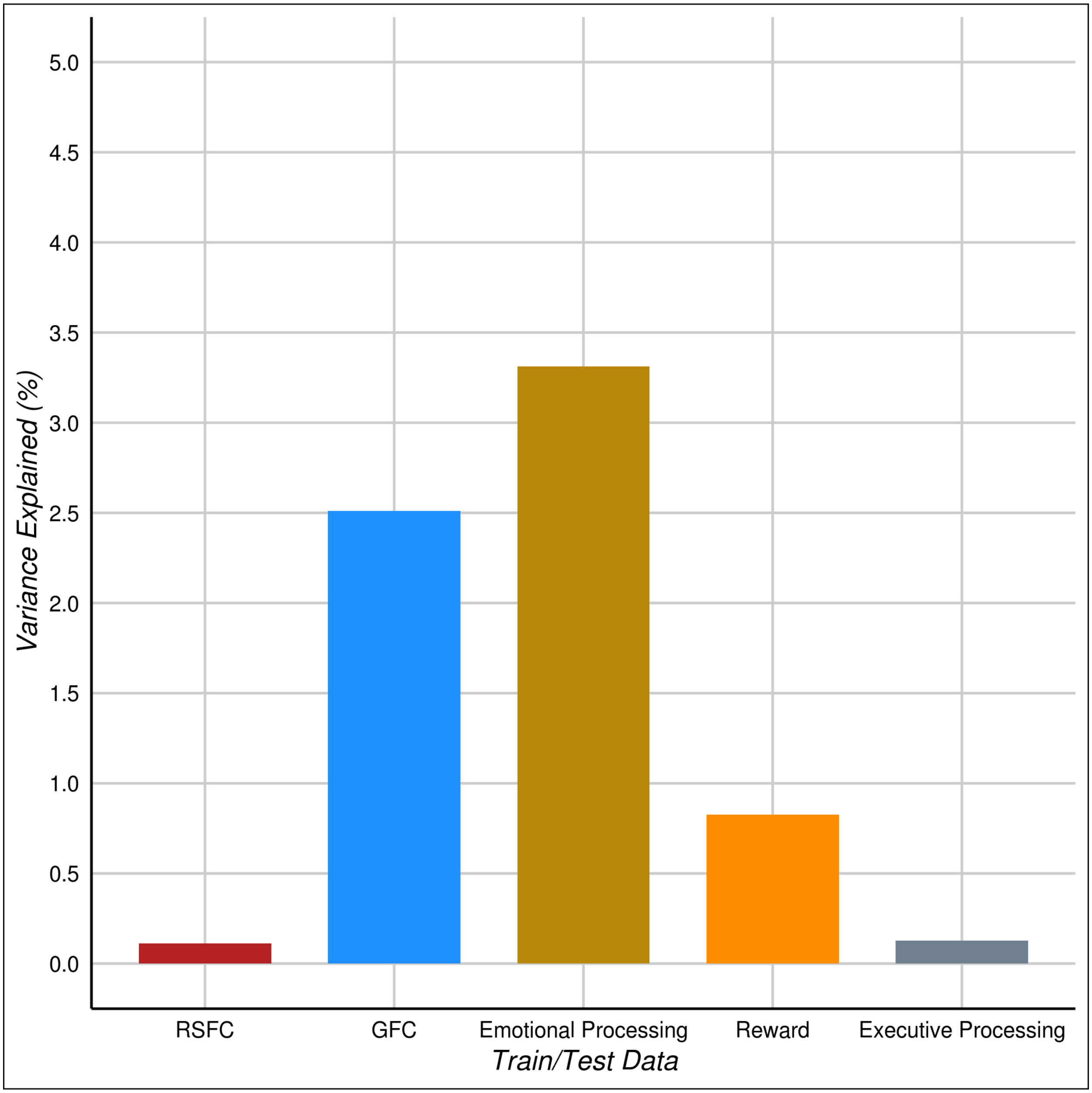

**Figure S6.** Out of sample predictions of cognitive ability from data trained in HCP and tested in Dunedin. While the general pattern is very similar to that of data trained in Dunedin and tested in HCP (Figure 6), the effect sizes are much smaller. These differences in out of sample prediction highlight the importance of training sample. Training in Dunedin likely leads to greater out of sample predictions because Dunedin is a larger sample, with a more reliable measure of cognitive ability. In addition, Dunedin is a population representative sample with a greater range in cognitive ability while HCP is a convenience sample that is highly-educated and range-restricted on cognitive ability.

| HCP ICCs | | |
|---|---|---|
| **Data Type** | **With Task Regression** | **Without Task Regression** |
| 10 mins GFC | 0.35 | 0.42 |
| 10 mins GFC w/o Rest | 0.35 | 0.43 |
| 10 mins Rest | 0.34 | 0.34 |
| 40 mins GFC | 0.56 | 0.58 |
| 40 mins GFC w/o Rest | 0.50 | 0.56 |
| 40 mins Rest | 0.53 | 0.53 |
| Emotion | 0.24 | 0.29 |
| Gambling | 0.23 | 0.35 |
| Language | 0.26 | 0.38 |
| Motor | 0.26 | 0.32 |
| Relational | 0.26 | 0.37 |
| Social | 0.30 | 0.40 |
| Working Memory | 0.28 | 0.40 |

**Table S1.** The influence of data composition and preprocessing method on average test-retest reliability (ICCs) of intrinsic connectivity. GFC constructed without rest has comparable ICCs to GFC constructed with rest. Task regression has an adverse effect on the ICCs of task-based data.

| Out of Sample Prediction | | |
|---|---|---|
| **Data Type** | **With Task Regression** | **Without Task Regression** |
| 10 mins GFC | 10.11 [4.37, 17.89] | 6.25 [1.93, 12.67] |
| 10 mins GFC w/o Rest | 12.74 [6.30, 21.44] | 9.61 [4.00, 17.56] |
| 40 mins GFC | 9.49 [4.28, 16.73] | 9.12 [3.96, 16.56] |
| 40 mins GFC w/o Rest | 13.99 [6.86, 22.28] | 2.86 [0.26, 7.67] |
| 40 mins RSFC | 5.66 [1.61, 11.63] | 5.66 [1.61, 11.63] |
| Emotion | 9.80 [4.33, 17.64] | 4.49 [1.10, 10.43] |
| Executive Function | 4.62 [1.02, 10.63] | 3.24 [0.42, 8.18] |
| Reward | 9.12 [3.84, 16.40] | 2.43 [0.20, 7.18] |

**Table S2.** The influence of data composition and preprocessing method on the predictive utility of cognitive ability from intrinsic connectivity. GFC constructed without rest has comparable ICCs to GFC constructed with rest. Task regression has an adverse effect on the ICCs of task-based data. All effect sizes are presented as % variance explained ($r^2$) with 95% confidence intervals in brackets.

**WITHIN SAMPLE PREDICTION (HCP)**

| FC DATA | Model | Correlation | p-value | # edges |
|---|---|---|---|---|
| **RSFC** | Positive | .22 | < .001 | 964 |
| **RSFC** | Negative | .18 | .002 | 858 |
| **RSFC** | Both | .20 | < .001 | 1822 |
| **GFC** | Positive | .30 | < .001 | 1710 |
| **GFC** | Negative | .28 | < .001 | 1684 |
| **GFC** | Both | .27 | < .001 | 3394 |

**Table S3.** Pearson correlations, p-values, and number of edges included in the predictions of cognitive ability. These values are from the within-sample predictions using the HCP dataset (Figure 5). Predictions are made from 40 minutes of RSFC or 40 minutes of GFC using CPM. All models are presented here (positive, negative, and both). Only the both model is reported in the main manuscript.

**WITHIN SAMPLE PREDICTION (DUNEDIN STUDY)**

| FC DATA | Model | Correlation | p-value | # edges |
|---|---|---|---|---|
| **RSFC** | Positive | .28 | < .001 | 968 |
| **RSFC** | Negative | .22 | < .001 | 934 |
| **RSFC** | Both | .26 | < .001 | 1902 |
| **GFC** | Positive | .29 | < .001 | 2186 |
| **GFC** | Negative | .30 | < .001 | 2156 |
| **GFC** | Both | .32 | < .001 | 4342 |

**Table S4.** Pearson correlations, p-values, and number of edges included in the predictions of cognitive ability. These values are from the within-sample predictions using the Dunedin Study dataset (Figure 5). Predictions are made from 40 minutes of RSFC or 40 minutes of GFC using CPM. All models are presented here (positive, negative, and both). Only the both model is reported in the main manuscript.

**OUT-OF-SAMPLE**
**(TRAIN: DUNEDIN STUDY, TEST: HCP)**

| FC DATA | Model | Correlation | p-value | # edges |
|---|---|---|---|---|
| **RSFC** | Positive | .26 | < .001 | 968 |
| **RSFC** | Negative | .12 | .03 | 932 |
| **RSFC** | Both | .24 | < .001 | 1900 |
| **GFC** | Positive | .29 | < .001 | 2184 |
| **GFC** | Negative | .26 | < .001 | 2152 |
| **GFC** | Both | .31 | < .001 | 4336 |
| **EMOTION** | Positive | .26 | < .001 | 1286 |
| **EMOTION** | Negative | .26 | < .001 | 1436 |
| **EMOTION** | Both | .31 | < .001 | 2722 |
| **REWARD** | Positive | .31 | < .001 | 940 |
| **REWARD** | Negative | .26 | < .001 | 798 |
| **REWARD** | Both | .30 | < .001 | 1738 |
| **EXECUTIVE** | Positive | .21 | < .001 | 1402 |
| **EXECUTIVE** | Negative | .23 | < .001 | 1454 |
| **EXECUTIVE** | Both | .22 | < .001 | 2856 |

**Table S5.** Pearson correlations, p-values, and number of edges included in out-of-sample predictions of cognitive ability displayed in Figures 5 and 6. All models were trained using the Dunedin Study dataset and tested in the HCP dataset using CPM. All models are presented here (positive, negative, and both). Only the both model is reported in the main manuscript.

| | Rest | Emotion | Gambling | Language | Motor | Relational | Social | Working Memory |
|---|---|---|---|---|---|---|---|---|
| Correlation With GFC | 0.98 | 0.94 | 0.98 | 0.93 | 0.94 | 0.92 | 0.93 | 0.93 |

**Table S6.** Correlations between the full intrinsic connectivity matrix from each task with 40 minutes of GFC in the HCP dataset. Correlations demonstrate that GFC captures common variance shared across task states rather than being driven by any single task.

| | Rest | Faces | Facename | Mid | Stroop |
|---|---|---|---|---|---|
| Correlation with GFC | 0.96 | 0.97 | 0.9 | 0.95 | 0.95 |

**Table S7.** Correlations between the full intrinsic connectivity matrix from each task with 40 minutes of GFC in the Dunedin Study dataset. Correlations demonstrate that GFC captures common variance shared across task states rather than being driven by any single task.

### Supplemental Experimental Procedures

#### Task fMRI paradigms

*Human Connectome Project.* Task fMRI paradigms for the Human Connectome Project are described extensively in (Barch et al., 2013).

*Dunedin Longitudinal Study*

*Emotion processing task.* The task consists of four blocks of a perceptual face-matching task interleaved with five blocks of a sensorimotor control task. The Dunedin Study version of this task consists of one block each of fearful, angry, surprised, and neutral facial expressions presented in a pseudorandom order across participants. During face-matching blocks, participants view a trio of faces and select one of two faces (on the bottom) identical to a target face (on the top). Each face processing block consists of six images, balanced for gender, all of which were derived from a standard set of pictures of facial affect. During the sensorimotor control blocks, participants view a trio of simple geometric shapes (circles and vertical and horizontal ellipses) and select one of two shapes (bottom) that are identical to a target shape (top). Each sensorimotor control block consists of six different shape trios. All blocks are preceded by a brief instruction ("Match Faces" or "Match Shapes") that lasts 2 s. In the task blocks, each of the six face trios is presented for 4 s with a variable interstimulus interval (ISI) of 2-6 s (mean = 4 s) for a total block length of 48 s. A variable ISI is used to minimize expectancy effects and resulting habituation and maximize amygdala reactivity throughout the paradigm. In the control blocks, each of the six shape trios is presented for 4 s with a fixed ISI of 2 s for a total block length of 36 s.

*Stroop task.* In this version of the Stroop task, participants identify the color of a target word in the center of a screen by selecting 1 of 4 identifier words. Selections are made by pressing 1 of 4 buttons on a button box, with each button matching an identifier word on the screen (e.g., index finger button 1 = identifier word on the far left, etc.). In congruent trials, targets were in colors congruent with the target words; in incongruent trials, targets are in colors incongruent with the targets. The four identifier words are in white in both conditions. After a 2 s fixation lead in, participants completed three 60 s blocks of congruent trials interleaved with three 60 s blocks of incongruent trials. Both conditions are followed by an 8 s fixation period for a total task time of 6 min and 50 s. Each block contains 12 trials, each consisting of 2 s stimuli, 1 s feedback, and a variable inter stimulus interval averaging 2 s.

*Monetary incentive delay (MID) task.* This version of an event-related MID task consists of 12 eight second trials for each of 3 conditions, presented in a pseudo-random order, for a total of 36 trials. Conditions consist of potential $1 reward, potential $5 reward, and no monetary outcome. On reward trials, participants could win money by pressing a button during the presentation of a target. During each trial, participants see either a green dollar amount cue (reward conditions) or a white “$0” (neutral condition) (cue, 2 seconds), then fixate on an “x” as they wait for a variable interval (delay, 2250–3000 ms), and then respond with a button press to a white target triangle that appears for a variable length of time (target, 70-680 ms) with a button press. Feedback (feedback, 2 seconds), which follows the disappearance of the target, notifies participants of whether they had successfully responded to the target with either the word “HIT” or “MISS”, and indicates their cumulative total at that point. Initial task difficulty (i.e. duration of the target) is based on reaction times collected during a practice session before scanning, and an adaptive algorithm was employed to adjust the target's duration such that each participant should succeed on ~66% of his or her target responses. Trials are separated by a variable (2-6 seconds) inter-trial interval. fMRI volume acquisitions are time-locked to the offset of each cue and thus were acquired during anticipatory delay periods.

*Memory encoding task.* This task consists of the encoding and subsequent recall of novel face-name pairs. A distractor task (odd/even number identification) is interleaved between encoding and recall blocks to prevent maintenance of information in working memory. During each of four encoding blocks, subjects view six novel face-name pairs for 3.5 seconds each. During each of four recall blocks, subjects view six faces each presented for 2 seconds and immediately followed by an incomplete name fragment for 1.5 seconds during which they are required by forced-choice to determine if the fragment is correct or incorrect. A 1.5 second inter-trial interval is used during recall blocks. During each of four distractor blocks, subjects view six different numbers for 3.5 seconds each and are required to determine if the numbers are odd or even.

**Task activation regression**

For the main analyses task activation was removed from the fMRI time series by including regressors processed with the canonical hemodynamic response function, corresponding to each task condition along with the other nuisance regressors in the 3dTproject step during pre-processing. Task regressors for each sample and task are described below.

*Human Connectome Project*. Regressors removed from the fMRI time series included 8 for the working memory task, corresponding to each combination of 0- or 2-back and four stimuli types (faces, body, places, tools); 3 for the gambling task, corresponding to reward, loss, and neutral trials; 6 for the motor task, corresponding to the blocks of movement for each of five body parts, and the cues across all conditions; 7 for the language task, corresponding to each combination of math or story condition and presentation, question, and response phase, as well as cues across all conditions; 2 for the social cognition task, corresponding to the mental interaction and random interaction blocks; up to 3 for the relational processing task, corresponding to relation and match blocks as well as error trials; and 2 for the emotional processing task, corresponding to blocks of faces and shapes.

*Dunedin Study.* Regressors removed from the fMRI time series included 5 for the emotion processing task, corresponding to blocks of shapes and faces with each of four different expressions; up to 4 for the Stroop task, corresponding to each combination of incongruent or congruent trials and correct or incorrect responses; up to 10 for the monetary incentive delay task, corresponding to the cue and anticipation period for the neutral, small gain, and large gain conditions, the target, and each combination of hit or miss feedback for each condition; and 3 regressors for the memory encoding task, corresponding to encoding, distractor, and recall blocks.

**Supplemental References**

Barch, D.M., Burgess, G.C., Harms, M.P., Petersen, S.E., Schlaggar, B.L., Corbetta, M., Glasser, M.F., Curtiss, S., Dixit, S., Feldt, C., Nolan, D., Bryant, E., Hartley, T., Footer, O., Bjork,

J.M., Poldrack, R., Smith, S., Johansen-Berg, H., Snyder, A.Z., Van Essen, D.C., 2013. Function in the human connectome: Task-fMRI and individual differences in behavior. Neuroimage 80, 169–189. https://doi.org/10.1016/j.neuroimage.2013.05.033
